# Expanding the utility of sequence comparisons using data from whole genomes

**DOI:** 10.1101/2020.01.15.908137

**Authors:** Sean Gosselin, Matthew S. Fullmer, Yutian Feng, Johann Peter Gogarten

## Abstract

Whole genome comparisons based on Average Nucleotide Identities (ANI), and the Genome-to-genome distance calculator have risen to prominence in rapidly classifying taxa using whole genome sequences. Some implementations have even been proposed as a new standard in species classification and have become a common technique for papers describing newly sequenced genomes. However, attempts to apply whole genome divergence data to delineation of higher taxonomic units, and to phylogenetic inference have had difficulty matching those produced by more complex phylogenetics methods. We present a novel method for generating reliable and statistically supported phylogenies using established ANI techniques. For the test cases to which we applied the developed approach we obtained accurate results up to at least the family level. The developed method uses non-parametric bootstrapping to gauge reliability of inferred groups. This method offers the opportunity make use of whole-genome comparison data that is already being generated to quickly produce accurate phylogenies. Additionally, the developed ANI methodology can assist classification of higher order taxonomic groups.

**Significance Statement:** The average nucleotide identity (ANI) measure and its iterations have come to dominate *in-silico* species delimitation in the past decade. Yet the problem of gene content has not been fully resolved, and attempts made to do so contain two metrics which makes interpretation difficult at times. We provide a new single based ANI metric created from the combination of genomic content and genomic identity measures. Our results show that this method can handle comparisons of genomes with divergent content or identity. Additionally, the metric can be used to create distance based phylogenetic trees that are comparable to other tree building methods, while also providing a tentative metric for categorizing organisms into higher level taxonomic classifications.

## Introduction

DNA-DNA Hybridization (DDH) holds the distinction of being the gold standard for species delineation (1). The method is technically challenging and its results at times are poorly reproducible across labs. Consequently, ongoing efforts attempt to supplement or replace DDH with *in silico* methods by taking advantage of the ongoing revolution in genome sequencing (2–6). One of the major approaches has been the Average Nucleotide Identity (ANI) (2).

ANI was first proposed in 2005. At the time the method used the average identity of shared open reading frames (ORFs) instead of the whole genome (2). The authors defined a species level ANI cutoff and examined large disparities in gene content among the strains and species in their dataset. A year later, they explored this metric in greater depth and observed that ANI was correlated with the percent of content shared, but that a significant amount of genomic nucleotide divergence (1-2%) needed to have occurred before there were major shifts in genome content (7). In 2007, the emphasis shifted from ORFs to the whole genome as the ANI method was adapted to directly compare to DDH (3). This shift led to the development of programs such as the jSpecies Java application which could perform the Goris method in a local and scalable manner (8). However, the consideration of the varying gene content became de-emphasized with the default exportable output from jSpecies not including any reference to shared content in comparisons. This de-emphasis on gene content is largely irrelevant when comparing closely related organisms due to the correlation between ANI and shared genome content. Yet this becomes a problem when only fractions of the genomes are shared, and can lead to spurious ANI results.

The problem of shared gene content was examined again in 2015 with the publication of the gANI method (6). This approach explicitly considers the shared gene content and offers two separate delimiters for a species: gANI (global ANI, which was based off the 2005 method), as well as an “Alignment Fraction” (AF), a measure of the proportion of genes shared. While gANI offers an important upgrade to the ANI paradigm it does contain an important limitation. Namely, there is no obvious answer on how to interpret a comparison between two taxa where the ANI is above the threshold and the AF is below, or vice-versa, which is a problem given that these metrics are most often used for species delimitation.

Here we suggest that ANI derived distance measures can also be used to reconstruct phylogenies that reflect shared ancestry, thus providing a natural extension to group species into genera and families. We introduce a single distance measure from whole genome data incorporating both the ANI and AF, labeled Total Average Nucleotide Identity (tANI) into the final metric. An advantage of the described method is that it can be applied to high quality draft genomes prior to annotation and gene clustering. Additional time is saved by using distance-based tree-building methods that are typically faster than maximum-likelihood or Bayesian inference methods. Ignoring phylogenetic information retained in individual gene families, this approach is not impacted by gene transfers that create misleading phylogenetic information – a gene acquired from outside the studied group will lower the alignment fraction, but it will not provide a signal moving the gene recipient closer to the root of the studied group. Furthermore, including the AF in the calculation of pairwise distances incorporates point mutations and gene transfers as processes of genome divergence into a single distance measure. We correct pair-wise distances for saturation and use bootstrap re-sampling to assess reliability. The analyzed test cases illustrate that this approach reliably resolves relations within genera and families.

## Results

### Necessity of Saturation Correction and AF incorporation

ANI values and the programs that calculate them were not designed with the intent of phylogenetic reconstruction. Consequently, the basic methodology works well within the confines of species delineation; however, the ANI values (or the corresponding sequence divergence) become prone to saturation and lose information when one attempts to compare more divergent taxa. To illustrate this, we took two of our datasets, the Rhodobacterales, and the Aeromonadales, (Table 1) (see Table S01 for detailed description of the datasets) and compared the ANI values calculated from JSpecies (8) to our tANI method (Fig. 1). As genome comparisons move away from the within species scale that ANI was designed for (2) the noise in the jSpecies ANI result become considerable. In extreme cases, the jSpecies ANI value for a comparison can border on the species cutoffs despite incorporating only a small fraction of the genomes. An example of this occurs in the Aeoromonadales dataset. *Aeromonas* bivalvium CECT7113T is found to have jSpecies ANI values around 94% when compared to *Aeromonas* media CECT4232T; however, the AF has a value of only 0.527 (significantly below the expected species cutoff). The effects of small alignment fraction and no correction for saturation is further illustrated in the topology of a distance tree inferred from uncorrected jSpecies ANI values (Fig. 2). These results from the Aeromonadales dataset clearly demonstrate the effect of saturation on phylogenetic reconstruction beyond the most closely related of taxa (9). Through incorporation of AF into the pairwise distance and correcting for saturation, the tANI method ameliorates the issues described above. Our distance values increase steadily while uncorrected jSpecies ANI enters the early stages of saturation at ~85% identity (Fig. 1). If using jSpecies ANI with the MUMmer algorithm, the saturation effects appear even earlier (data not shown). We want to emphasize that our comparison with uncorrected ANI values should not be seen as a criticism of the original ANI methods, rather we use the comparison to illustrate the importance of considering AF and saturation in case ANI is used to infer shared ancestry.

**Table 1.** A bridged Dataset Descriptions

| Dataset<sup>*</sup> | Composition<sup>†</sup> | Remarks |
| --- | --- | --- |
| Aeromonas | Drawn from Colston, Fullmer et al. 2014. Only has *Aeromonas* genomes. | Chosen as these genomes already had MLSA and core genome phylogenetic trees constructed, allowing for us to more easily compare our method to these. |
| Aeromonadales | Composed of several *Aeromonas* genomes, the remaining available Aeromonadales outside of the *Aeromonas*, and several Enterobacterales which served as an outgroup. | With genomes from two separate orders, this dataset provides opportunity to explore outer limits of the method in regards to taxonomic range. |
| Rhodobacterales | Consists of Rhodobacterales genomes with a leaning towards the genera *Leisingera*, *Loktanella*, and *Ruegeria*. | Chosen for familiarity and for previously made MLSA for a subset of the taxa which provided an easy route for expansion. |
| Frankiales | Consists of a selection of publicly available Frankiales genomes. | Selected for the variety of genome sizes and GC content present within the Order, allowing us to check for biases. |

<sup>\*</sup>Based on dominant taxa in the dataset <sup>†</sup>A more comprehensive breakdown of the dataset is available in supplemental materials.

**Fig. 1.**
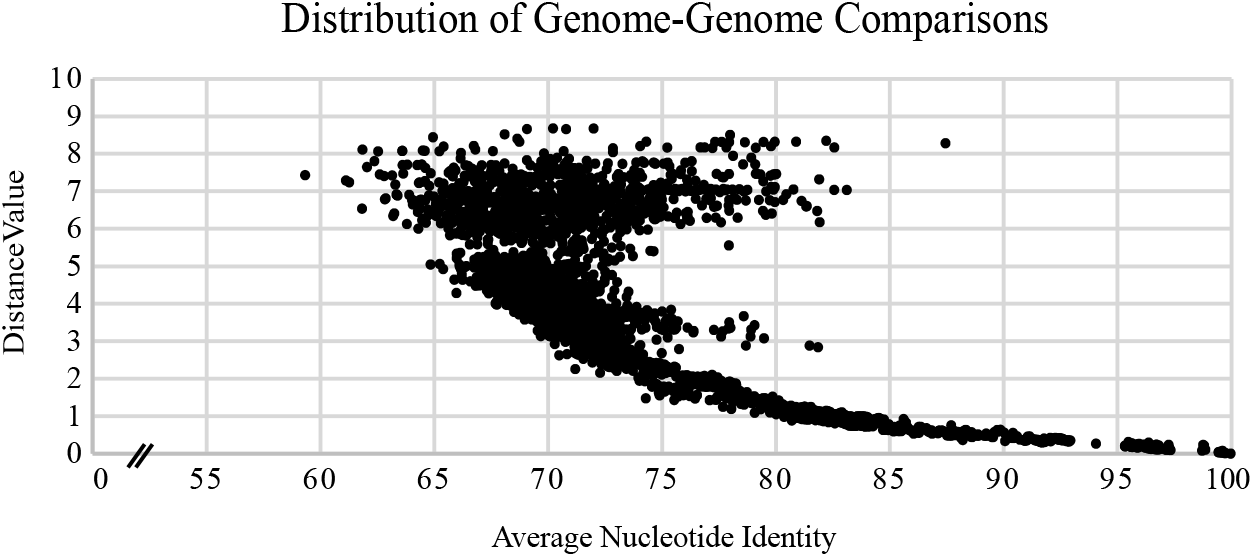
tANI distance value as a function of uncorrected jSpecies ANI value. This plot comprises individual genome-genome comparisons from both the Aeromonadales and Rhodobacterales datasets; resulting in a dataset of 6195 comparisons. This “tornado” configuration illustrates how jSpecies ANI begins to enter saturation by approximately 87%. This saturation is a function of declining AF values and sequence saturation.

**Fig. 2.**
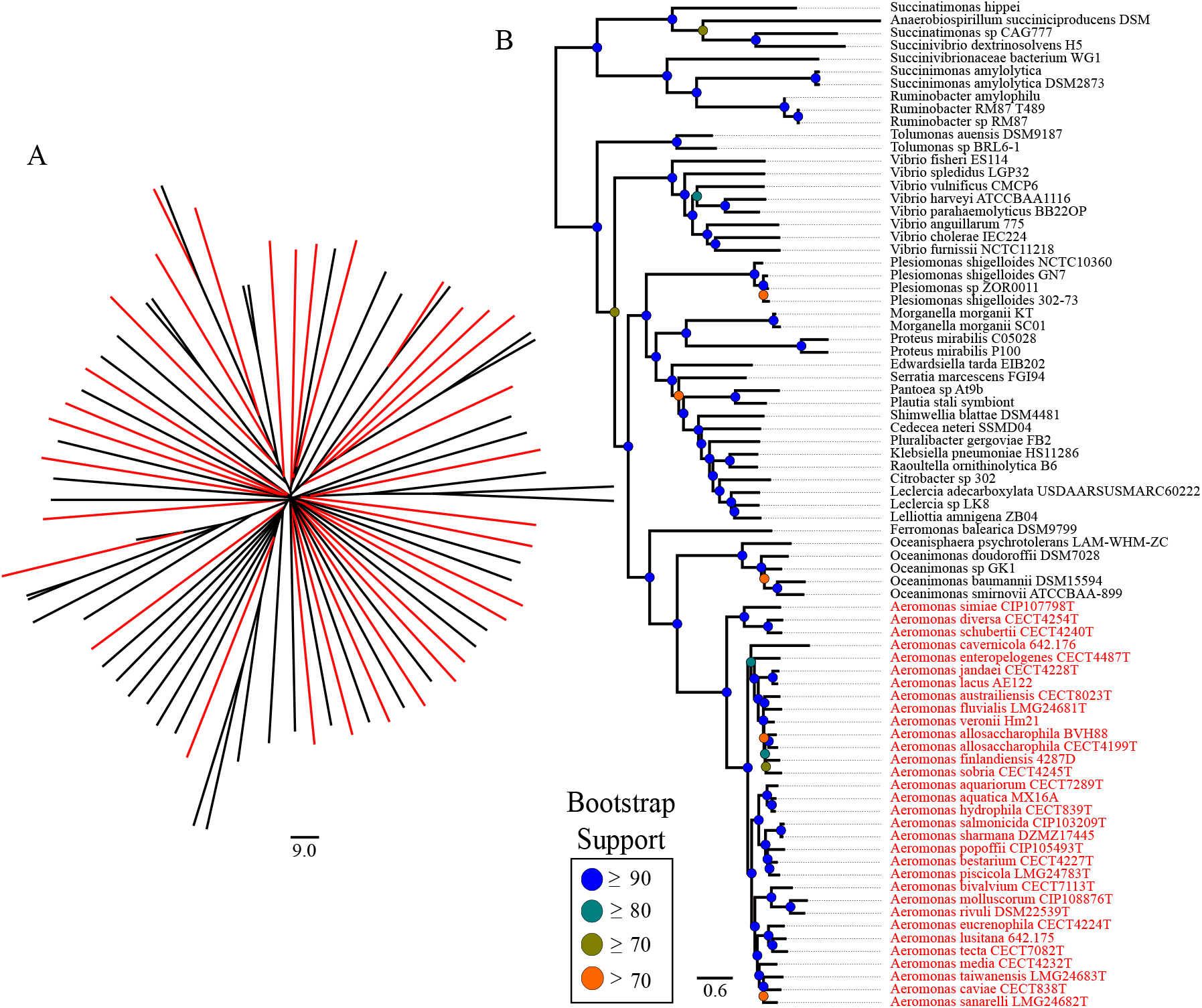
FastME phylogenies of the same Aeromonadales dataset. (A) The phylogeny inferred from jSpecies ANI values, converted to uncorrected distances (100-%ANI). The color-coding on A reflects the location of members from the *Aeromonas* genus within the dataset. (B) The phylogeny on the right is created from the same dataset using the tANI methodology before using FastME. All *Aeromonas* members are highlighted in red to illustrate their placement as a single clade.

### Genome Size and GC Content Do Not Create a Detectable Bias

Since our distance measure is based on the whole genome, differences in genomic traits, such as size and GC-content, could bias the results of the calculations and introduce artefacts into the final phylogeny and their support values. To test this, we developed a dataset using the order *Frankiales* (Table 1), composed primarily of the genus *Frankia*. This group has high variance in genome size (~4Mb to ~11 Mb) and considerable range of GC-contents (~60% to ~75%) which made it an ideal test case.

The tANI based distance tree for the *Frankiales* (Fig. 3B) set was very similar to the MLSA-derived reference phylogeny (Fig. 3A) (see the “Accuracy of the tANI Methods Compared to Multi-Gene Methods” section for a more detailed analysis). Mapping the size of the genome onto the tANI phylogeny showed no pattern of clustering by genome size (Fig. S1 A). While some groups cluster with similar sizes (e.g. the *F. coriariae* and *F. alni* clades), they match the MLSA topology and do not consistently group with only similar sized genomes. Mapping the GC-content onto the tANI phylogeny produced a similar result (Fig. S1 B), with no obvious patterns of GC-content bias.

**Fig. 3.**
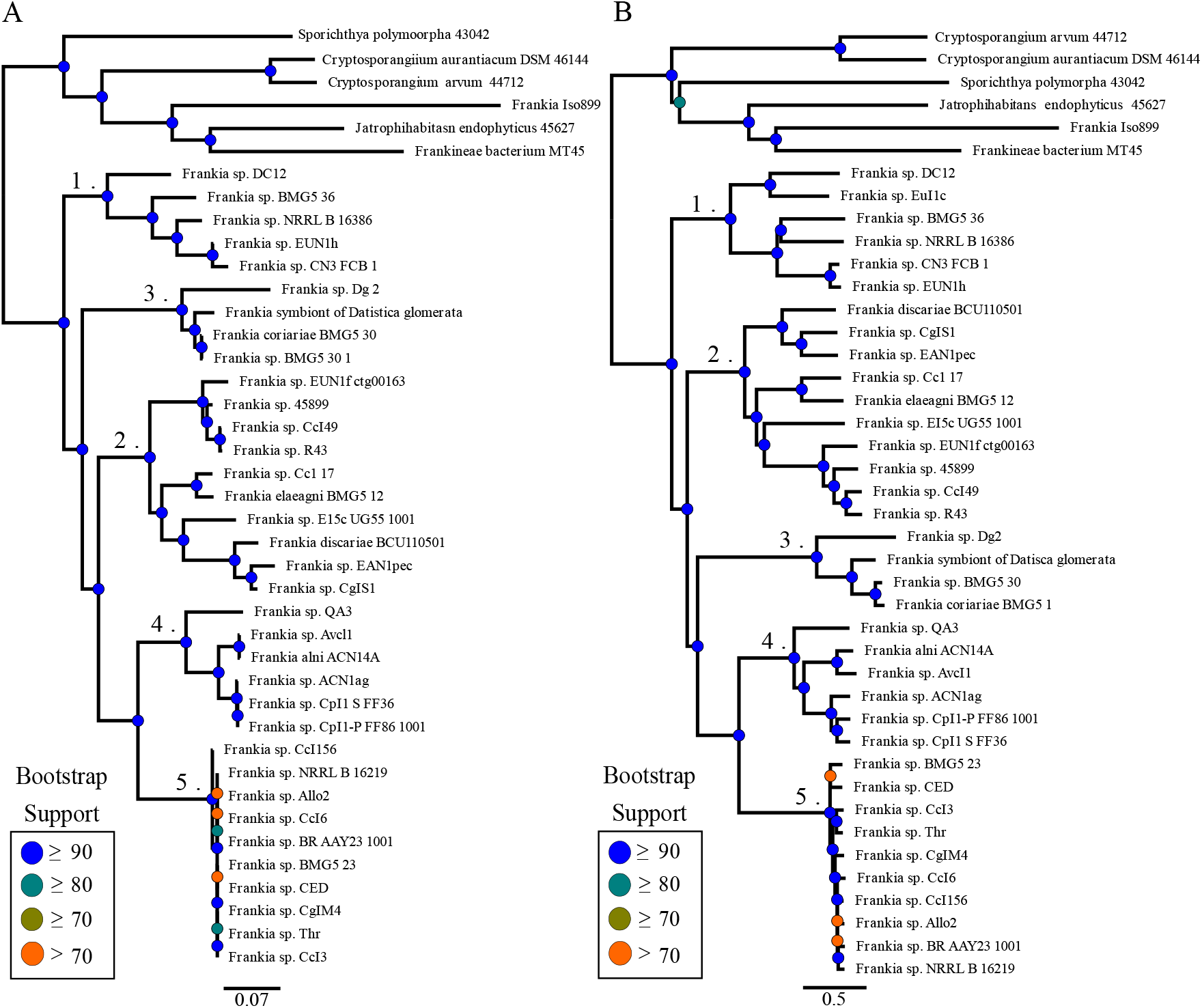
Phylogenies of the Frankiales dataset using two different methodologies. (A) The tANI derived phylogeny (left) compared against (B) the MLSA phylogeny (left). See Materials and Methods for details on the methods used phylogenetic reconstruction.

### Bootstrap Confidence Sets for tANI and core genome ML analyses are similar

To provide support values for our distance-based phylogenies, our script creates a set of non-parametric bootstrapped distance matrices (see material and methods for details). Inter-node certainty (IC) scores were calculated to assess the statistical uncertainty of the trees derived from non-parametric bootstrapping (in the following labeled as “support sets”). IC scores were calculated by mapping statistical support sets against reference trees (the tree derived from the original data without bootstrapping) as implemented in RAxML v8.1 (10, 11). IC represents a quantification of the level of disagreement in a support set for a particular node in a phylogeny; a higher score indicates less disagreement between topologies. The tree certainty average (TCA) value is the average of IC values across the entire tree, representing an assessment of overall conflict between the support set and reference phylogeny (10). The Aeromonas dataset (Table 1) was used as a test case as it offers an expanded core phylogeny in addition to the MLSA, allowing a comparison between different whole genome methods. Comparing support datasets against the best tree calculated using the same method, the TCA for the Aeromonas tANI phylogeny was 0.86, 0.87 for the expanded core genome phylogeny, and .65 for the MLSA phylogeny. Comparing between approaches (MLSA, tANI, Mashtree) results in positive TCAs, *i.e.* the trees agree with one another more than they disagree; however, the scores are below 0.4, with the exception of tANI and core genomes based analyses for the Aeromonas dataset, which resulted in TCAs of 0.61 (Table S2). To further compare our bootstrap method to other approaches, we calculated the Robinson-Foulds distances within each of the support sets from the MLSA and tANI method and analyzed the pairwise distances using principal coordinate analysis (PCoA) (Fig. 4). The PCoA plot shows that the statistical support sets for both of the datasets co-localize with one another; indicating that the support sets from the different methods represent similar tree space.

**Fig. 4.**
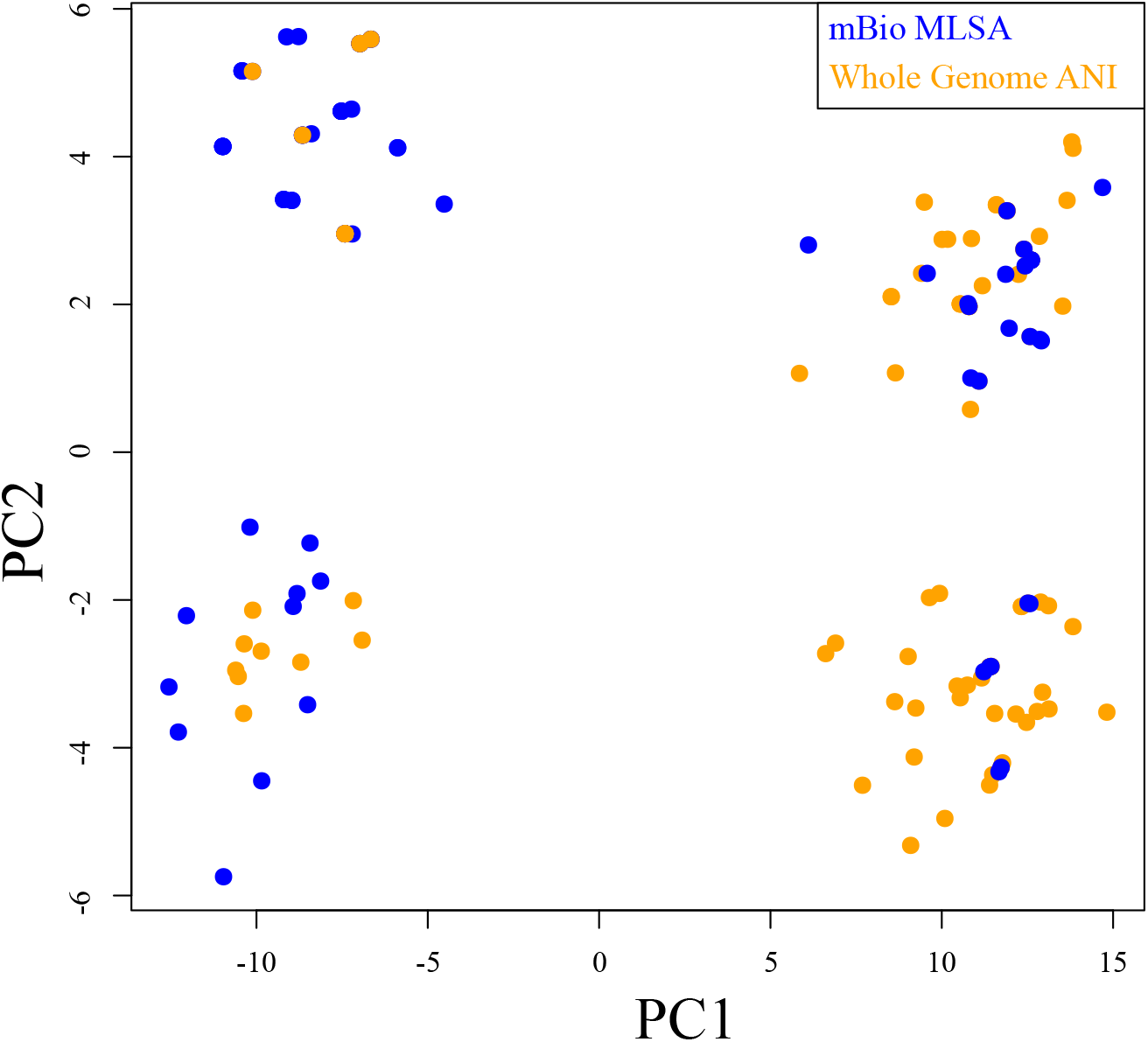
PCoA plot of the distance between trees from bootstrap samples calculated from the *Aeromonas* core genome, and tANI methods. The sample from the tANI method are colored in orange (Whole Genome ANI) and the *Aeromonas* core genome bootstraps are in blue (mBio MLSA). The support sets overlap in every cluster, suggesting that the two methods capture similar topologies.

### Accuracy of the tANI Methods Compared to Multi-Gene Methods

For the *Aeromonas* test dataset, differences between the extended core phylogeny and the tANI derived phylogeny are the placements of *Aeromonas veronii* AMC34 and the *A. allosaccharophila* clade (Fig. 5). *Aeromonas veronii* AMC34 is still placed within the extended A. *veronii, sobria* and *allosaccharophila* clade using the tANI method, but tANI disagrees on the specific location, and places AMC34 at the base of the *veronii* group, instead of base of the entire clade. This placement at the base of the *veronii* group shifts the A. *allosaccharophila* and A. *sobria* strains to a more basal position in this clade. However, AMC34’s placement is poorly supported in the tANI based analysis. Deeper clades within the tANI phylogeny match those of the extended core phylogeny, and are highly supported.

**Fig. 5.**
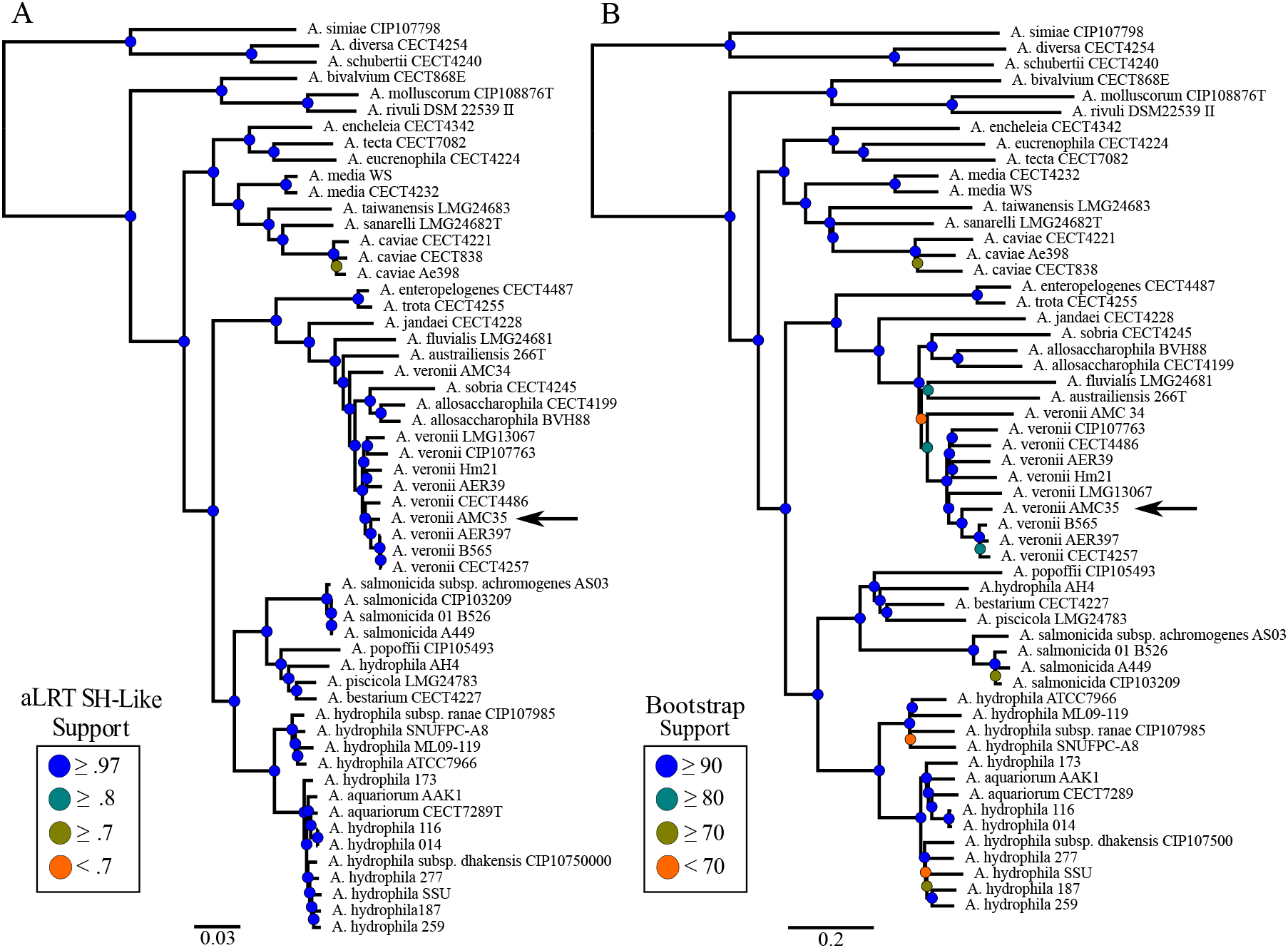
Comparison of *Aeromonas* phylogenies reconstructed using different methodologies. (A) The Extended Core Phylogeny, inferred using Approximate Maximum-likelihood (Colston et al., 2014), and (B) the tANI methodology, inferred using Fast Minimum Evolution. Keys for node support apply to the tree directly to the right of the key. Arrows point to the location of *Aeromonas veronii* AMC35.

The phylogenies produced by the extended core MLSA and tANI for the Frankiales dataset also had few differences (Fig. 3). Principle of the these was the divergence of clades 2 and 3 since tANI and MLSA disagree over whether clade 2 or clade 3 split off first. There is further disagreement on the placement of individual taxa primarily within clade 2 (see the placement of *Frankia* sp. Discariae BCU110501, *Frankia* sp. CgIs1, and *Frankia* sp. EAN1pec). However, aside from these minor disagreements within the clades, and differing levels of confidence (see bootstrap support, especially within clade 5), these two methods largely reproduce the same phylogeny.

Comparing different methods for the Rhodobacterales dataset yielded more complex results (Fig. S2). Both the tANI and the core genome MLSA phylogenies have low levels of support for the internal branches of most of the phylogeny, and disagree on the placement of the genera *Ruegeria, Loktanella, Roseobacter*, and the several other singletons. The tANI and MLSA trees also disagree on the placement of *Loktanella*, with the tANI grouping it as one small paraphyletic clade and several smaller monophyletic clades, whereas the MLSA method groups *Loktanella* into one large paraphyletic group and one small monphyletic group. tANI and MLSA trees both agree on keeping the Caulobacterales a monophyletic clade, and keep the *Leisingera* genus together, both with high support. Within *Leisingera* there is minor disagreement on the placement of the individual strains, but they are largely kept in the same branching pattern. The *Ruegeria* groupings are also kept intact across the two trees. Further comparison of the tANI method on the Rhodobacterales dataset against other methods (see Mashtree section below) implies the large amount of internal disagreement is implicit to the dataset, and will require more detailed analysis to untangle.

Additional visual confirmation for the results described above is provided by the split graphs created for each of the datasets (see supplemental figures 3,4,5 and 6).

### Comparison of tANI Method with Mashtree

Genome-based phylogenies have been in the literature for some time. As such, it is appropriate to compare our methodology with other available whole-genome methods and assess our methodologies strengths and weaknesses. To this end we compare our method to Mashtree (https://github.com/lskatz/mashtree), which is an extension of the Mash kmer-calculation (12)(Ondov et al., 2016).

For the Aeromonas dataset, Mashtree had only minor disagreements with our method (Fig. S7 A). For example, MashTree moved the placement of *A. media*, and shuffled members within the A. *salmonicida* and A. *aquatica* groups. This pattern generally repeats itself in the Rhodobacterales dataset (Fig. S7 B). Mashtree also kept *Leisingera, Rhodobacter* and the major *Ruegeria* clade together in a similar fashion to the tANI phylogeny. Additionally, the MashTree phylogeny generally agrees with the branching patterns the tANI phylogeny proposes, while deviating at nodes of low support in the tANI and MSLA based phylogenies. However, Mashtree did separate *Loktanella* into a number of monophyletic clades. Comparing the Mashtree topology with support sets from tANI, MLSA, and core genome analyses gave results similar to the other TCA values comparing between methods (Table S02).

### This Novel Extension of ANI Matches Older Methodologies

Since tANI is based off a species delimitation intended measure, we wanted to see if it maintained this original purpose while also being able to produce phylogenies. To determine the species cutoff based on a single genome-to-genome distance calculation we used a receiver operating characteristic curve (ROC) analysis. Working on the union of the Aeromonas and Rhodobacterales datasets, the ROC estimates a distance cutoff of 0.315, at a specificity of 99.984 and sensitivity of 99.200 against the accepted nomenclature (Fig. 6A). Examination of the ROCs for the constituent datasets reveals that the two genera are not equally easy to classify (Fig. 6B, 6C). However, when taxa in the Rhodobacterales set are reclas-sified along the lines suggested in the MLSA phylogeny (see supplemental table 3), the genera curve improves in sensitivity from 80% to 99% while maintaining the same specificity (Fig. 6D).

**Fig. 6.**
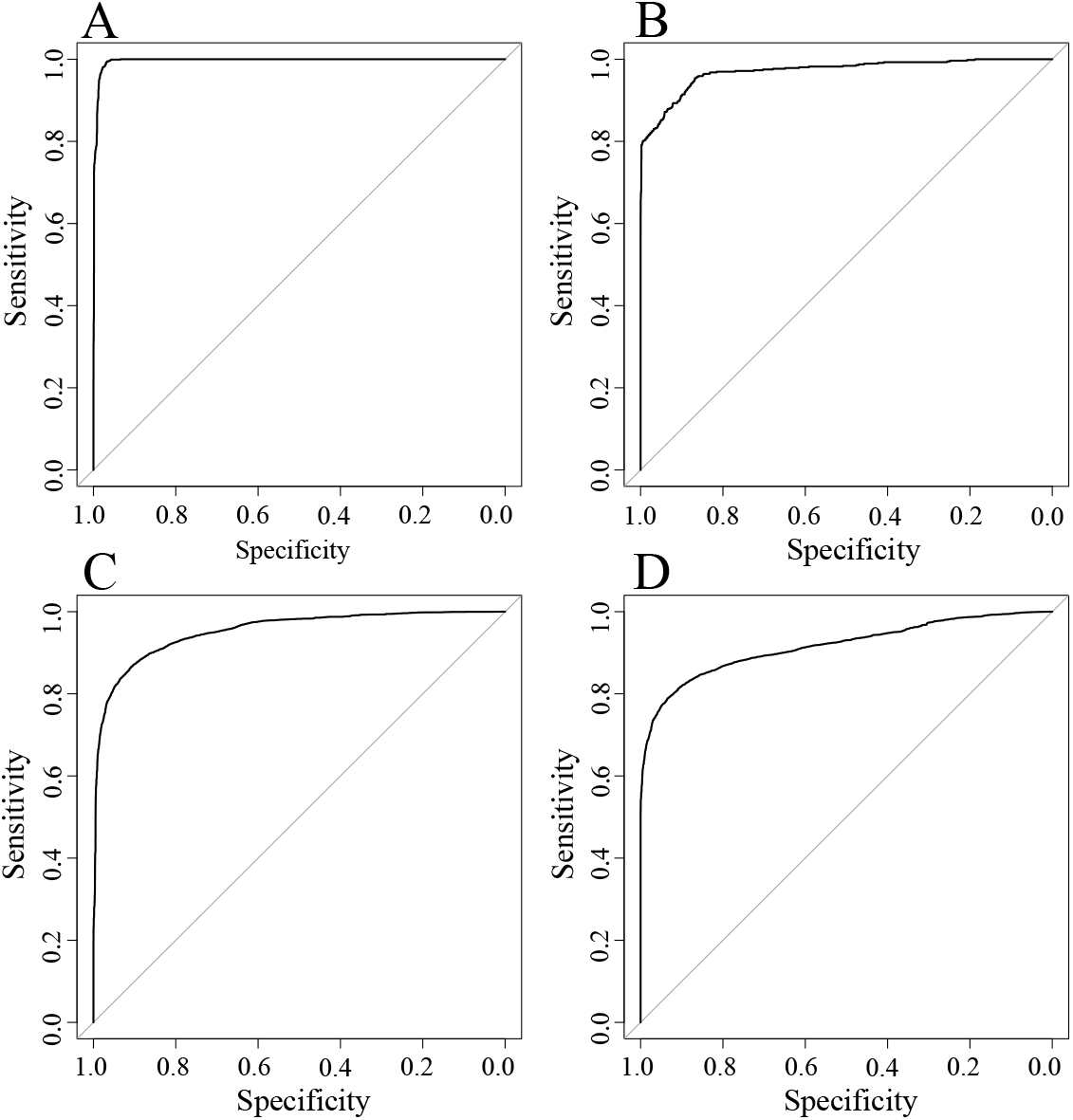
Response Operator Curves reporting the sensitivity and specificity of the tANI at discriminating species relationships. Plots show (A) the union of the Aeromonas and Rhodobacterales datasets against accepted nomenclature (specificity of 99.98%, and sensitivity of 99.20%), (B) the Aeromonas dataset (specificity of 96.68%, and sensitivity of 97.97%), (C) the Rhodobacterales dataset (specificity of 83.78%, and sensitivity of 80.09%), (D) the Rhodobacterales dataset after reclassifying taxa (specificity of 83.31%, and sensitivity of 99.13%).

### Novel tANI Method Offers the Ability to Delimit Deeper Taxonomic Ranks

One added benefit from our use of broader taxonomic samplings in some of our datasets is the opportunity to test our distance measure against ANI and GGDC species cutoffs. When the distances for every pair-wise comparison from the Aeromonadales and the Rhodobacterales sets were plotted, and filtered for taxa suspected of misclassification (see supplemental figure 8 for a version using NCBI classifications), a series of recognizable peaks for each taxonomic rank were observed (Fig. 7). The ROCs were used to provide statistical evidence for these observations. At the genus level, the Aeromonadales (Fig. 8) and Rhodobacterales sets (Fig. 8B) have similar distance cutoffs (3.3 and 3.4, respectively) and varied but generally high specificities (96.7% and 83.3%) and sensitivities (98.0% and 99.1%). At the family level, the combined datasets returned a cutoff of 4.57 and maintained specificity of 90.7% and sensitivity of 86.5% (Fig. 8C). At the order level, the combined datasets fell off to 4.42 cutoff, 94.2% specificity and 71.44% sensitivity, suggesting the method could not discriminate at this taxonomic rank (Fig. 8D). It should be noted, that these values are likely to be highly dataset specific.

**Fig. 7.**
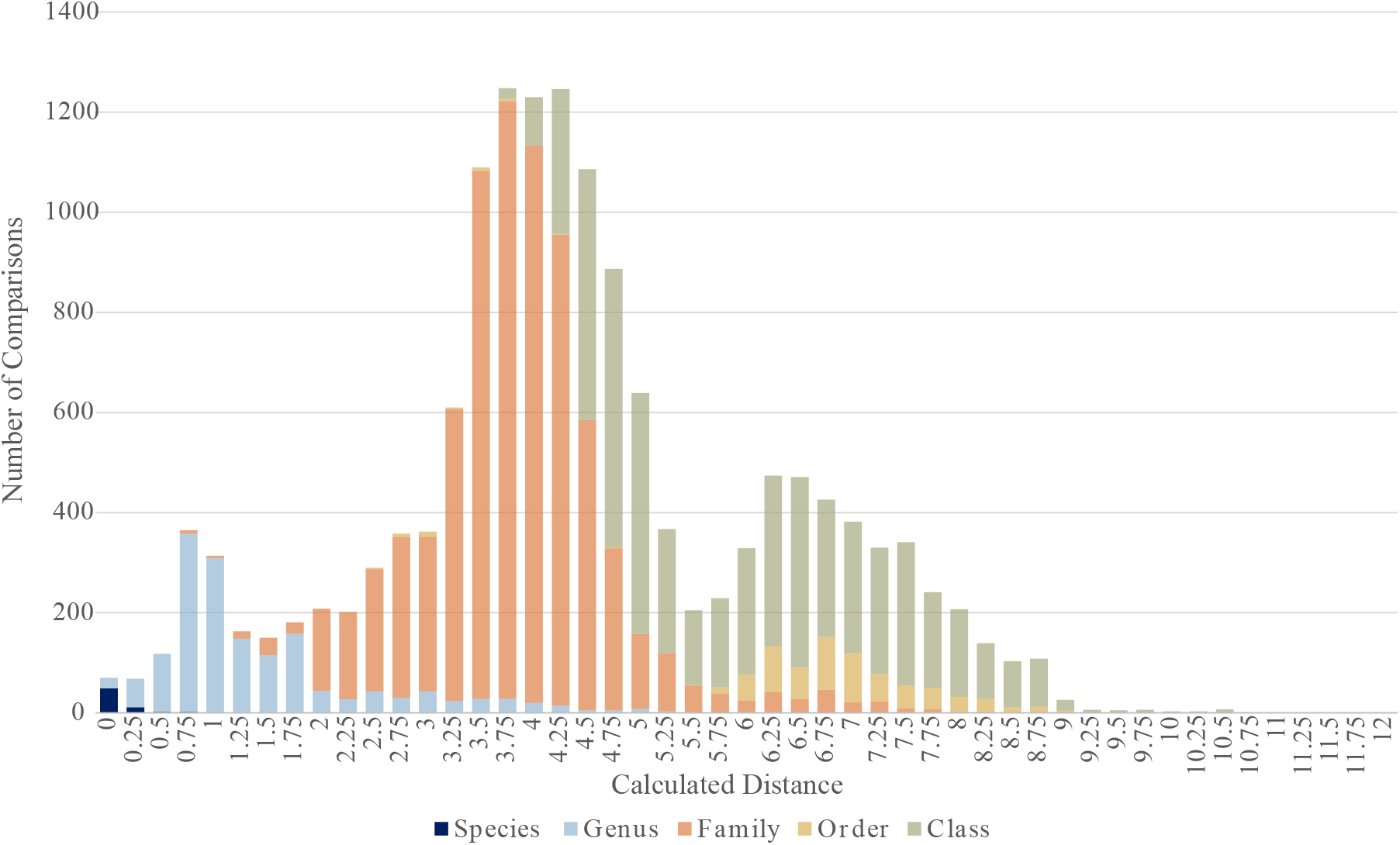
Histograms of tANI values for taxonomic rank comparisons in our datasets using re-categorization of misclassified taxa along the lines suggested in the reference phylogenies. A figure using the NCBI taxonomy classification scheme is included as supplementary figure S3

**Fig. 8.**
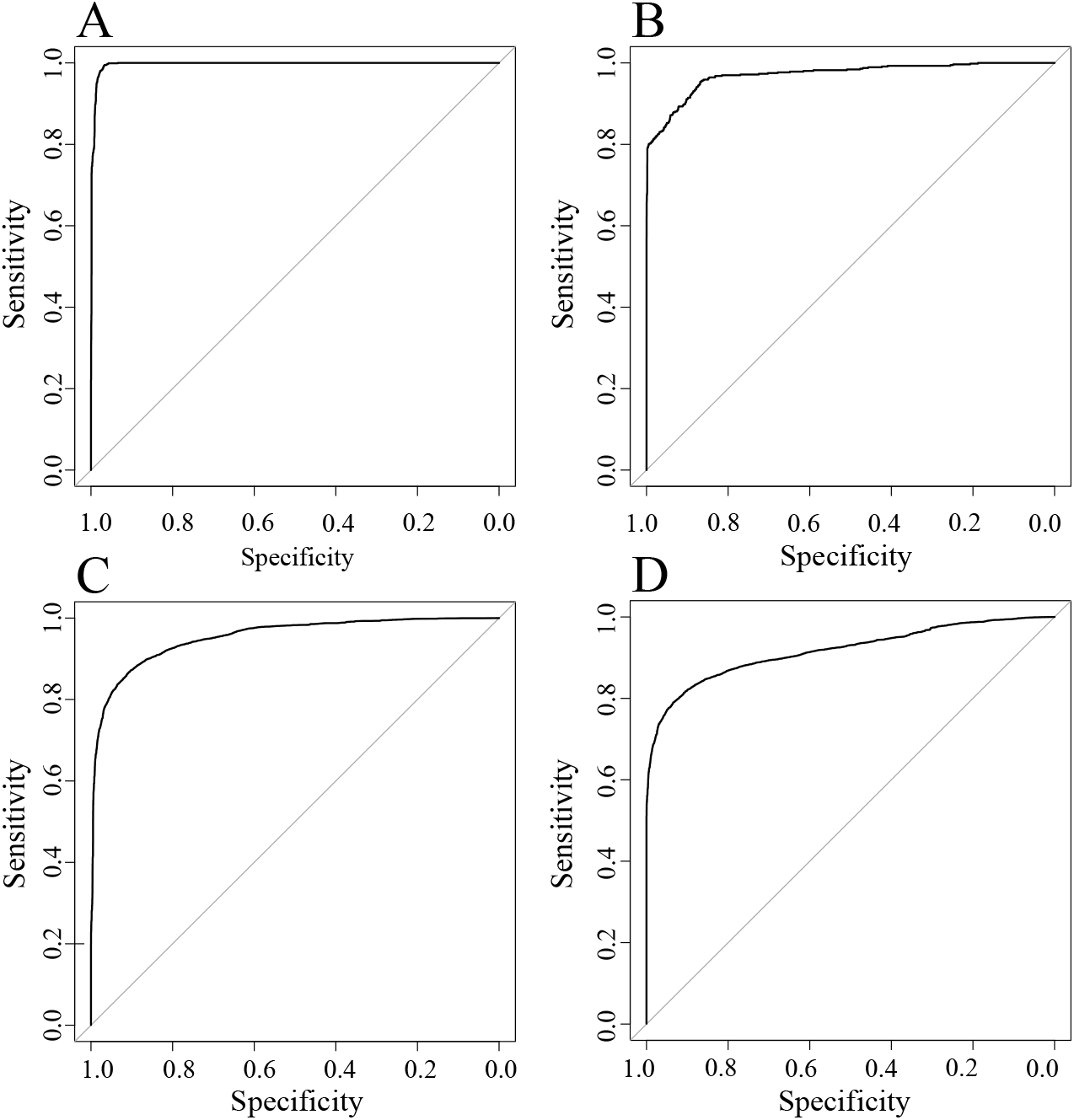
Response operator curves reporting the sensitivity and specificity of the ANI-distance at discriminating deeper taxonomic relationships for (A) the AeroOG dataset at the genus level, (B) the Roseo dataset, also at the genus level. Specificities (96.7% and 83.3%) and sensitivities (98.0% and 99.1%) are varied but generally high. Panel C shows our combined datasets at the family level. The family relationships maintain an ability to discriminate between classifications at rate close to the genus data (90.7% specificity and sensitivity of 86.5%). Panel D displays the combined data at order level. While order level specificity was high (94.2%) its sensitivity was only 71.4%, suggesting the method is breaking down and losing the ability to discriminate.

### Misclassified Taxa

A number of taxa in our datasets appear to be misclassified under incorrect genera, family, order, and species labels (Table S3). These taxa fall into groups for which phylogenetic analyses support their misclassification. The ROC determined cutoffs also supported that these taxa are outside of their assigned group. These taxa were reclassified into novel groups along their phylogenetic lines for the purpose of our taxonomic rank cut-off analyses (Table S4). Our tANI metric cutoff agreed with these decisions, and when redoing the ROC analyses with these changes improved the sensitivities and specificities of those cutoffs. There are three specific higher order classifications to which this applies: Loktanella, Ruegeria, and Succinivibrionaceae. Additionally, several species level classifications may need to be revised, specifically those mentioned in Table S3.

## Discussion

### Success of tree-building

The tANI method has demonstrated the capacity to match more sophisticated techniques. tANI trees consistently showed comparable levels of conflict to reference phylogenies and matched the level of confidence displayed by other methods such as MLSA when examining the datasets used within this paper. The tANI methodology performed well at the species, genus, family, and order levels; the relationships observed in the reference trees held true in the our tANI trees. Furthermore, TCA tests have shown that our bootstrapping methodology shares a significant portion of the uncertainty that other support methods provide (Fig. 4). These phylogenies, and associated tests have provided evidence to demonstrate the suitability for using ANI to infer phylogenies to at least the order level and likely into higher ranks. The implemented bootstrap support values provide a means to assess if genomes that are too divergent are included in an analysis.

### tANI is not overwhelmed by biases

The core of this work is predicated on the assumption that the genome as a whole conveys a significant amount of relevant information about the history of the organism. This assumption is broadly comparable to those made in using genomic content information to infer phylogeny and is subject to many of the same critiques (13). There are two primary issues to consider.

First, in light of potentially rampant horizontal gene transfer (HGT), how much of a cell’s genome will reflect a history of cell divisions rather than a composite of signals from the organism’s recombination partners? Fortunately, in many instances HGT and shared ancestry reinforce one another (14, 15). How much this applies to deeper taxonomic ranks, however, is, unfortunately, not certain. It is possible that the flows of gene-sharing that unite and divide such close relatives as *Escherichia* and *Salmonella* may not behave in the same way with more distant relationships. For deep divergences a genome-based approach may fail because of highways of gene sharing (16); however, regarding the evolution within orders, gene transfer can be considered as one process contributing to the gradual divergence of genomes (14) and contributes to tANI based distances. This gradual divergence is reflected in a smaller alignment fraction in case of transfers that add a new gene to the recipient genome, and in decreased nucleotide identity in case the transfer results in replacement of a homologous. Our analyses of the Frankiales genomes (Fig. S1) show that even in case of large differences in genome size due to deletion, duplication, and gene transfer the tANI based genome distances capture the same phylogenetic signal that is retained in genes that are present in all the analyzed genomes.

In general, the tANI based approaches for within-order phylogenies compare well with those obtained through genome core and MLSA analyses. The extent to which the noted differences reflect lower resolution and certainty for the tANI based distances in between genera comparisons, or the stronger impact of gene transfer events on sequence-based methods remains to be determined. Different combinations of core genes can strongly support contradicting phylogenies (17), suggesting that phylogenies from concatenated aligned sequences should not automatically be considered more reliable.

### Misclassified taxa

Results from our methods on the Rhodobacterales dataset show that there is a clear separation of the *Loktanella* and *Ruegeria* genera into multiple separate clades; however, *Lok-tanella* is significantly more fragmented (Fig. S2). The conclusion that these classifications should be re-described is supported by results from previous literature on *Ruegeria.* While some studies supported a monophyletic clade (18, 19), these studies lack many of the strains and taxa currently available, and the consistent non-monophyletic nature observed in our study has been duplicated in other recent studies with similar species sampling (9, 20). *Loktanella* may also require a revisit, as previous literature would suggest that our results (Fig. S2) are more reflective of the actual phylogeny. Newer studies of the genus and the larger groups to which they belong have included higher taxon sampling in their phylogenetic analyses, which provide support for this non-monphyletic interpretation of the genus (9, 20).

The *Aeromonadales* dataset suggests that the higher order classification of *Succinivibrionaceae* within the order may also be up for reconsideration (Fig. 2). Members of the family *Succinivibrionaceae* are extremely distant from the rest of the *Aeromonadales* order, with distance values reaching saturation. These values are so large that they commonly dwarfed the distance values calculated between other members of *Aeromonadales* and the distant members of Gammaproteobacteria and *Enterobacteriaceae.* The individual *Succinivibrionaceae* may be grouping together as the result of long branch attraction, though it is difficult to assess the family in higher detail, as there are few sequences publicly available. In addition, the original classification of *Aeromonadales* did not include the family *Succinivibrionaceae* (21) and no further analyses were reported that confirmed they should be included. This classification was seemingly the result of one 16S study (22) and no further phylogenetic analyses appears to back this claim.

### Deeper taxonomic ranks

In the same sense that ANI and GGDC have been used to delimit species (and in the case of GGDC strains), we examined if tANI distances could provide a first indication to discriminate between genus, family and order relationships. Clearly, grouping in higher taxonomic levels should be based on phylogenetic analyses; however, distance values can provide a first indication, especially in cases of poor taxon sampling. While our test sets are not exhaustive, the results were promising. Using an optimal cut-off level (as determined using criteria determined by Youden 1950) genus assignments were achieved at a rate of ~10% false positives and false negatives at ~1%. At family level, the false positives remained roughly unchanged, but the false negatives increased to ~14%. As with previous iterations of ANI, different groups will require specific considerations outside of a one cutoff fits all mold, as is evident given slight variances in optimal cutoffs for the different datasets.

### Conclusion

We have identified a valuable extension to the comparative analysis of whole-genome data that are being routinely generated by researchers. The ability to produce viable and statistically supported phylogenies in this manner offers the possibility for researchers to save time on what would otherwise be more complex and time-consuming phylogenomic techniques. For within family analyses, the phylogenies generated via the tANI method are robust and match the confidence and accuracy of current popular techniques and other whole-genome metrics. The discrimination power of the tANI method falls off when different families from the same order are includes. Furthermore, the possibility that the tANI method can provide preliminary evidence to help differentiate deeper taxonomic relationships offers the potential that it may be able to assist or provide evidence in favor of classification schemes going forward. Finally, many researchers are already producing information that is key to the described methodology, and can be easily transitioned for use in the tANI method. The tANI distance-based method and sequence-based methods (MLSA and core gene concatenations) have different sensitivity towards artifacts created through gene transfer from outside the group under analysis. We recommend inclusion of tANI based phylogenies as one of the tools to infer within family relationships.

## Methods

### Genomes used

The genomes used in this study are either draft whole genome assemblies or complete assemblies available via NCBI (Reference, Table S1). Selection initially centered on two groups for which previous phylogenetic and phylogenomic work had been done by this group. The first, the Aeromonas dataset, encompasses the 56 *Aeromonas* genomes used in Colston et al. (2014) and represents a genus level taxonomic unit. The second, the Rhodobacterales dataset, encompasses those used in Collins et al. (2015) and Gromek et al. (2016) plus additions to investigate the cases of *Loktanella* and *Ruegeria* (9, 23). This set corresponds closely to a family level taxonomic unit (exempting the genera: *Phenylobacterium, Parvularcula, Maricaulis, Hyphomonas, Hirschia, Caulobacter, Brevundimonas*, and *Asticcacaulis*, which are used as outgroups to root the phylogeny). A third set, aimed at encompassing a broader phylogenetic and taxonomic range was created by adding all publically available non-*Aeromonas* Aeromonadales genomes to a subset of the Aeromonas dataset along with taxa outside the order including members of the Enterobacteriales. All together this group is called the referred to simply as the Aeromonadales dataset. As the name implies, this set corresponds to an order level unit. Finally, the available genomes from the order Frankiales were formed into another dataset (of the same name) with the intention to test the robustness of the tANI method to heterogeneous genome sizes and GC-contents.

### Reference Phylogenies

Comparison reference phylogenies were obtained or generated for each dataset. For Aeromonas, the MLSA and expanded core phylogenies were obtained from Colston et al. (2014) (5). A reference for the Rhodobacterales dataset was generated by replicating the method described in Collins et al. (2015) (9), but with added *Loktanella* and *Ruegeria* genomes from NCBI. The Aeromonadales was calculated following the MLSA methodology described in Colston et al. (2014) for the included genomes (5).

The Frankiales reference required the *de novo* creation of an MLSA scheme in the absence of thorough examples in the literature. Twenty-four single-copy housekeeping genes were selected to form the alignment (Table S5). Nucleotide sequences for each gene were retrieved via BLAST from *Frankia casuarinae* (Accession: NC_007777.1) (reference for BLAST). BLASTn (v2.6.0) (24) was executed with the gene sequences as the query and the genomes as the target sequence. The coding sequences corresponding to highest scoring hits (using e-values) for each gene in a singular genome were aligned and concatenated. This was repeated for every genome, generating the multi-locus sequence alignment (MLSA) file. IQTree (v1.5.5) was executed with the MLSA file and built the phylogenetic tree with 1000 ultrafast-bootstraps (25–28). IQTree’s model finder arrived at the SCHN05 empirical codon model with empirical codon frequencies (+F) and Free Rate (29) model of rate heterogeneity with nine categories (+R5).

### ANI and AF Calculation

ANI is calculated in a similar methodology to that described by Varghese et al. (2015) such that ANI is not simply the sum of best hit identities over the total number of genes, but is instead described by the formula: ANI=∑(ID%*Length of Alignment)/(∑ (Length of the shorter fragment). Alignment fraction is described as: AF=∑(Length of the shorter fragment)/(Length of the Query Genome). The ID%, Length of the Alignment, and Length of the shorter fragment terms refer to the individual blast hits from genome-genome comparisons (see below). Our methodology differs from Varghese et al. (2015) in two respects. First, we do not limit our search to open reading frames but rather use the full scaffold/contig set of an organism. Second, we fracture the genomes into 1,020 nucleotide fragments in line with previous iterations of ANI calculation (2, 8). The fragments from the query genome were each compared to the reference genome via BLAST+ (v2.7.1). Results were filtered based on coverage and percent identity values and only the top bidirectional best hit was retained per segment. Filtered results were used to calculate the ANI and AF as defined earlier. The distance (abbreviated Total Average Nucleotide Identity, or tANI) was calculated by using the formula: **tANI =−log(AF*ANI)**. The natural log added to this calculation ensures that higher distance values correlate with genomes that have a lower ANI or AF (i.e. more dissimilar).

### Bootstrap Replicates

After genomes were split into 1,020 nucleotide segments, individual segments were chosen randomly with replacement from the pool of all of said genomes segments and used to create a new dataset. This new dataset was then compared against all other using the tANI methodology to create a bootstrapped distance matrix. These matrices were then used to infer their own trees. Those trees was then mapped onto the best tree to provide node support.

### Coverage and Percent Identity Cutoffs

The original percent identity and coverage cutoff values were chosen based on those laid down by Varghese et al. (6). Cutoff values were tested within the *Aeromonas* dataset. Average distance within the clade was measured over a range of cutoff values (Fig. S9) and multiple potential cutoffs were tested against the jSpecies ANI standard cutoffs of 70% identity and 70% length. We tested various cutoff values’ ability to construct phylogenetic trees compared to more conventional methods and concluded that 70-at-70 still produced the most accurate trees.

### Phylogenies from Distances

Tree-building from distance matrices was accomplished using the R packages Ape and Phangorn (30, 31). The balanced minimum evolution algorithm as implemented in the FastME function of APE was used to generate phylogenies for each distance matrix (32). Parameters used were: nni = TRUE, spr= TRUE, tbr = TRUE. A “best tree” was calculated from the point estimate values (original DDH estimations in *is*DDH; the initial calculated distance matrix in tANI) and a collection of bootstrap topologies from the resampled matrices. Support values were mapped onto the best tree using the function *plotBS* in Phangorn (31).

Split graphs were constructed from the distance matrices using Splitstree4 (33). Graphs were built using a NeighborNet distance transformation, ordinary least squares variance, and a lambda fraction of 1.

### Bootstrap evaluation

Tree certainty scores were calculated using the IC/TC score calculation algorithm implemented in RAxML v8 (10, 11). Tree distances were calculated using the R package Ape (32) and the treedist function of Phangorn (31).

### Residual Operating Characteristic Curve Analysis

A residual operating characteristic (ROC) curve was used to determine the optimal species cutoff for a single genome-to-genome distance calculation. Genomes from the sets of Aeromonadales and Rhodobacterales listed in the genome table were compiled and matrices of both the distance and raw jSpecies ANI were compiled from the set. The jSpecies ANI values were used to delimit species from the genomes selected. Each comparison was assigned a 1 if the comparison met the species cutoff, and a 0 if it did not according to jSpecies cutoffs (8). This list of 1’s and 0’s represents the true state.

True states and distance values were then compiled into a two-column data set. The R package pROC (35) allowed us to create a curve from the data and then determine the best cutoff values for the given set of data such that true negatives and true positives based on the cutoff value were maximized using methodology previously described (36).

## Acknowledgements

This work was supported in part through grants from the NSF (#1616514, PI, Mukul Bansal) and #1716046 to JPG). We thank Artemis Louyakis for many discussions and for critically reading drafts of the manuscript, and the Computational Biology Core in UConn’s Institute for Systems Genomics for providing computational resources.

## Supplemental Material

**Fig. S1** Phylogenies of the Frankiales dataset built from the tANI methodology and color coded to investigate potential biases of the method. (A) plots the length of the genome on each tip, and (B) plots the GC content of the genome on the tip. Neither A or B shows biased patterns.

**Fig. S2** Comparison of phylogenies reconstructed from the Rhodobacterales dataset. (A) The multi-gene phylogeny from Collins et al., (2015), and (B) the tANI distance based phylogeny. Both sides use bootstrapped node support on a scale from 100-0 indicated by the key.

**Fig. S3** Splits graph diagram for the Rhodobacterales dataset built using Splitstree4. The tANI method served provided the distance matrix used to build the graph. The graph reveals the part of this dataset that deviates from a tree-like description.

**Fig. S4** Splits graph diagram for the Frankiales dataset built using Splitstree4. The tANI method served provided the distance matrix used to build the graph. The graph supports splitting the Frankiales into 5 major groups, and one outgroup (Cryptosporangium, Sporichthya, Jatrophihabitans, etc).

**Fig. S5** Splits graph diagram for the Aeromonas dataset built using Splitstree4. The tANI method served provided the distance matrix used to build the graph. The graph backs previous understanding that the Aeromonas group partake in a large amount of horizontal gene-transfer.

**Fig. S6** Splits graph diagram for the Aeromonadales dataset built using Splitstree4. The tANI method served provided the distance matrix used to build the graph. The graph further backs the notion that the Aeromonadacea likely need to be revised, as the Succinivibrionaceae place further from the Aeromonas than those of the Enterobacteria.

**Fig. S7** Phylogenies derived from another whole-genome method, MashTree (37). (A) is constructed from the Aeromonas dataset, and (B) is built from the Rhodobacterales dataset.

**Fig. S8** Histograms of tANI values for taxonomic rank comparisons in our datasets using the uncorrected taxonomy as derived from NCBI. Compare Fig. 7 for a similar histogram using the data after correction for misclassified taxa.

**Fig. S9** Plot of the average distance of genome-genome tANI calculations within the *Aeromonas* dataset using varied percent identity and coverage cutoffs. This provided context to which regions of the plot would be best while building phylogenies. Specifically those cutoffs at which the distance was not saturated as a result of including low quality information (10-60), and not inflating as a result of filtering too much information (80+).

**Table S1** Full dataset description.

**Table S2** Average IC values

**Table S3** Changes for species level cutoff comparison

**Table S4** Suggested reclassification

**Table S5** Frankiales MLSA genes

