## Supplemental for "Expanding the utility of sequence comparisons using data from whole genomes"

**This PDF file includes:**

Figures S1 to S9
Tables S1 to S5

**Figures**

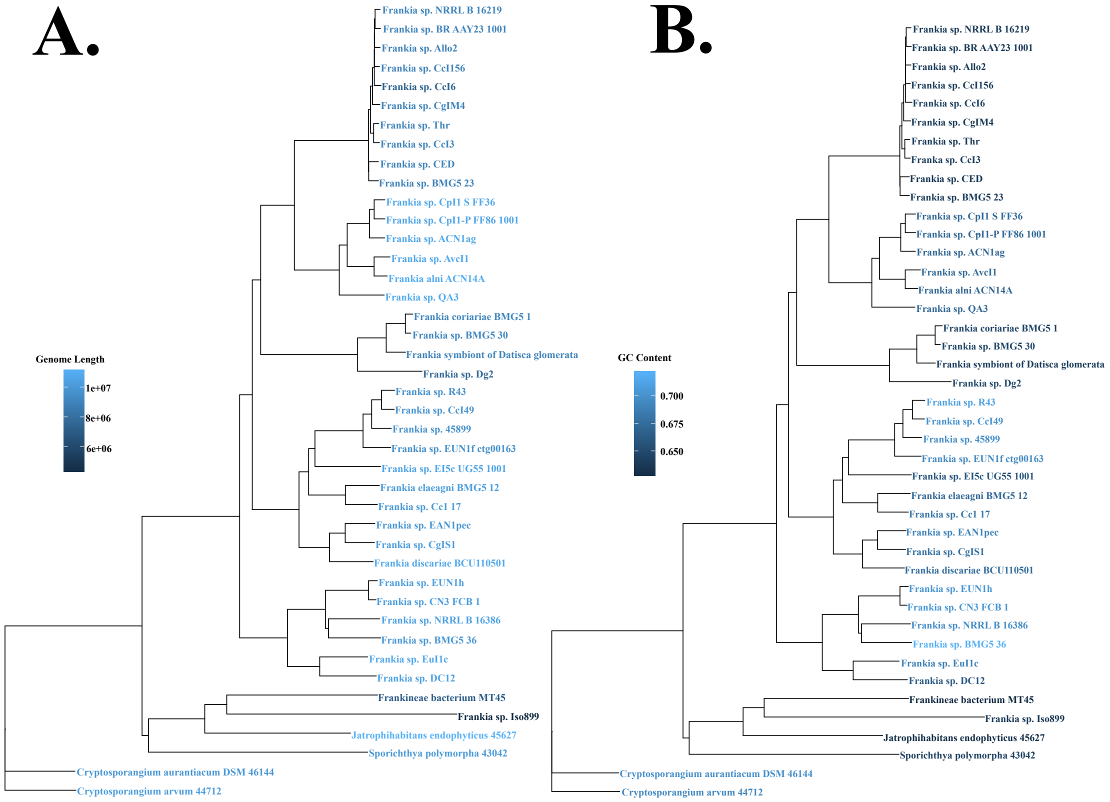


**Fig. S1.** Phylogenies of the Frankiales dataset built from the tANI methodology and color coded to investigate potential biases of the method. (A) plots the length of the genome on each tip, and (B) plots the GC content of the genome on the tip. Neither A or B shows biased patterns.

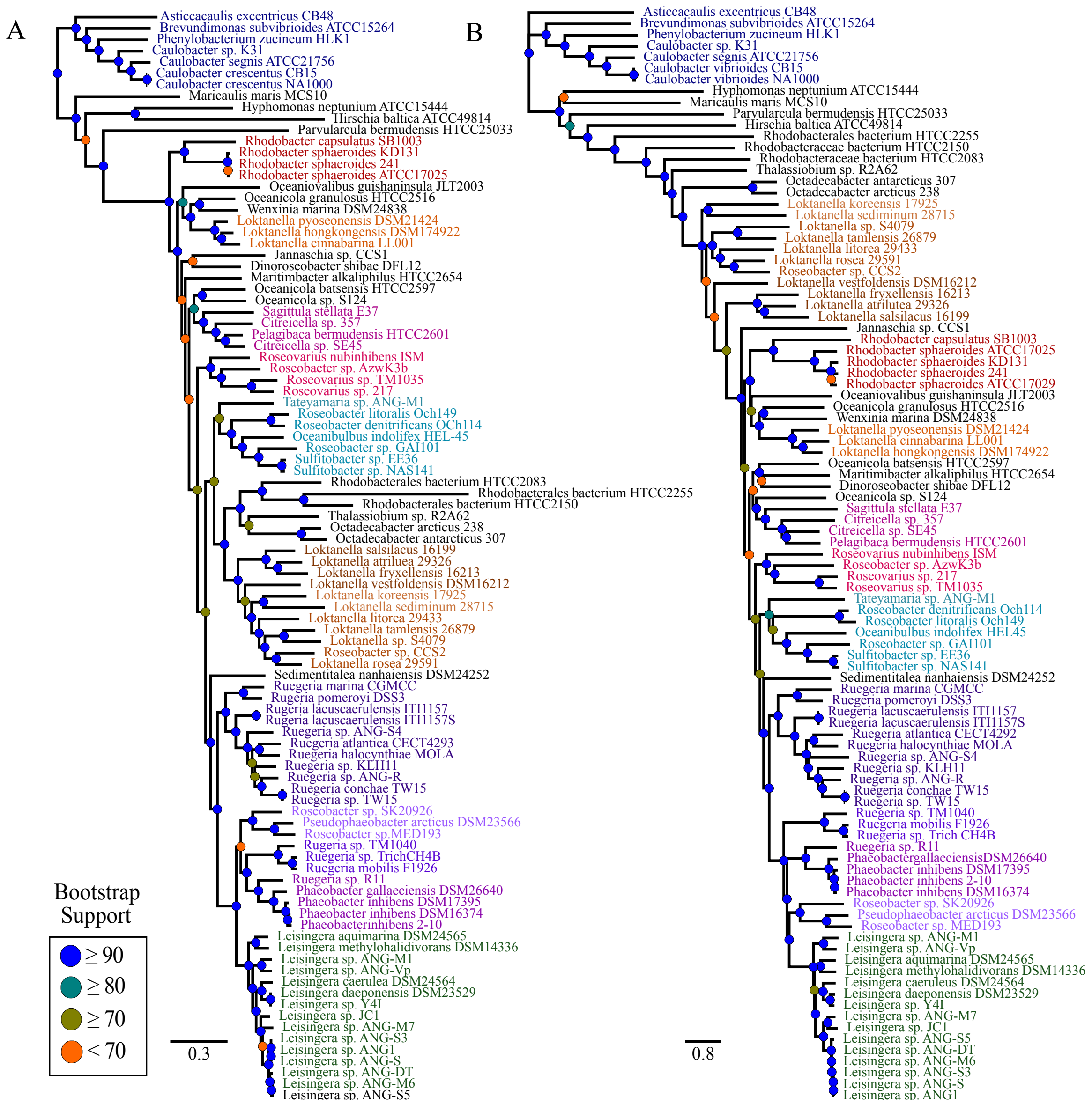


**Fig. S2.** Comparison of phylogenies reconstructed from the Rhodobacterales dataset. (A) The multi-gene phylogeny from Collins et al., (2015), and (B) the tANI distance based phylogeny. Both sides use bootstrapped node support on a scale from 100-0 indicated by the key.

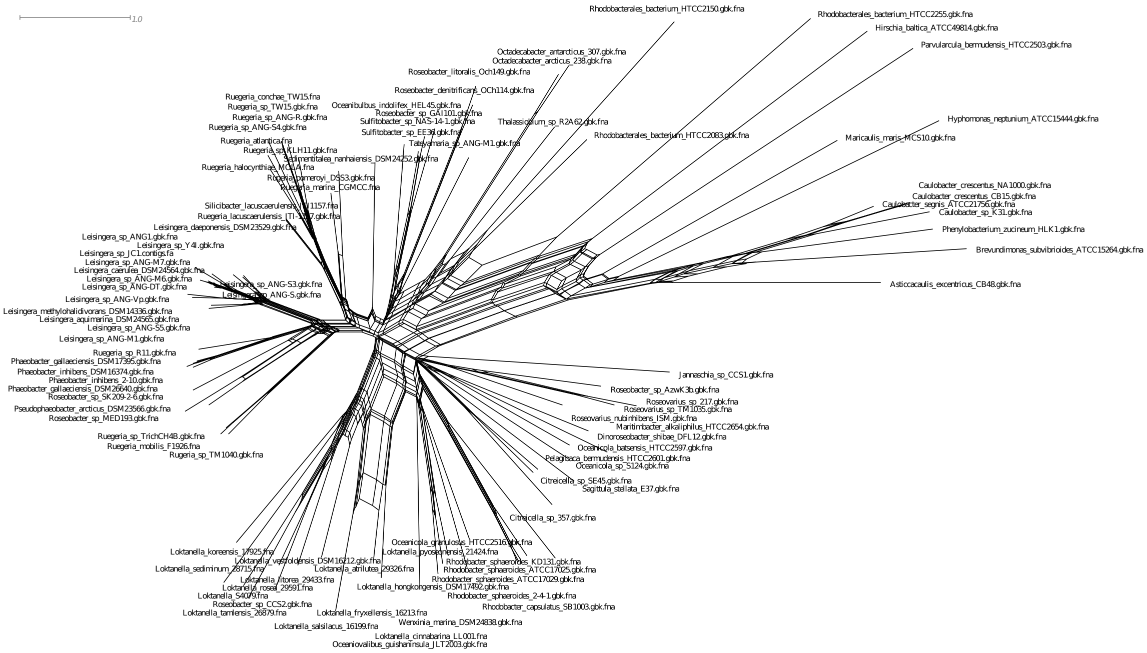


**Fig. S3.** Splits graph diagram for the Rhodobacterales dataset built using Splitstree4. The tANI method served provided the distance matrix used to build the graph. The graph reveals the part of this dataset that deviates from a tree-like description.

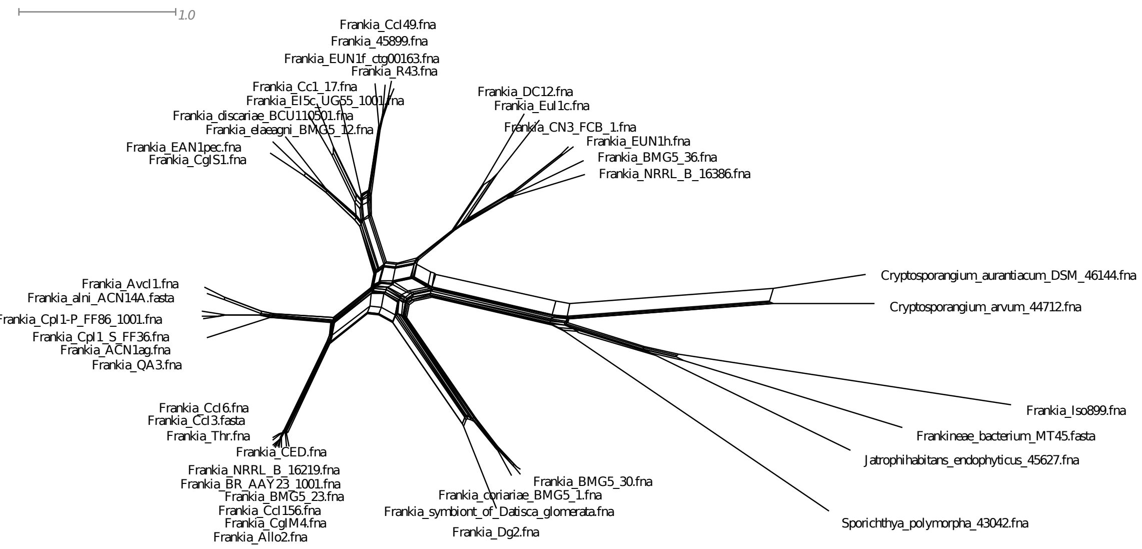


**Fig. S4.** Splits graph diagram for the Frankiales dataset built using Splitstree4. The tANI method served provided the distance matrix used to build the graph. The graph supports splitting the Frankiales into 5 major groups, and one outgroup (Cryptosporangium, Sporichthya, Jatrophihabitans, etc).

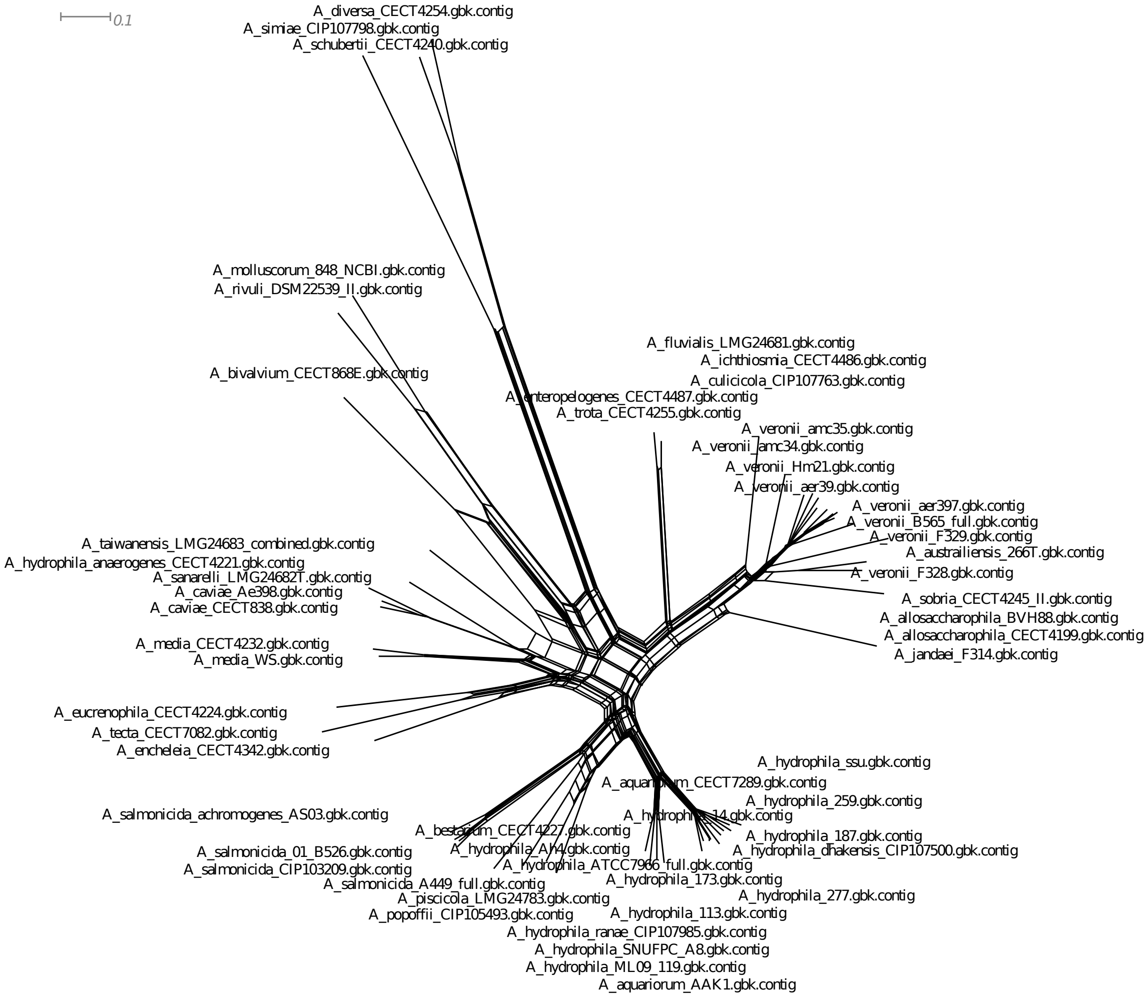


**Fig. S5.** Splits graph diagram for the Aeromonas dataset built using Splitstree4. The tANI method served provided the distance matrix used to build the graph. The graph backs previous understanding that the Aeromonas group partake in a large amount of horizontal gene-transfer.

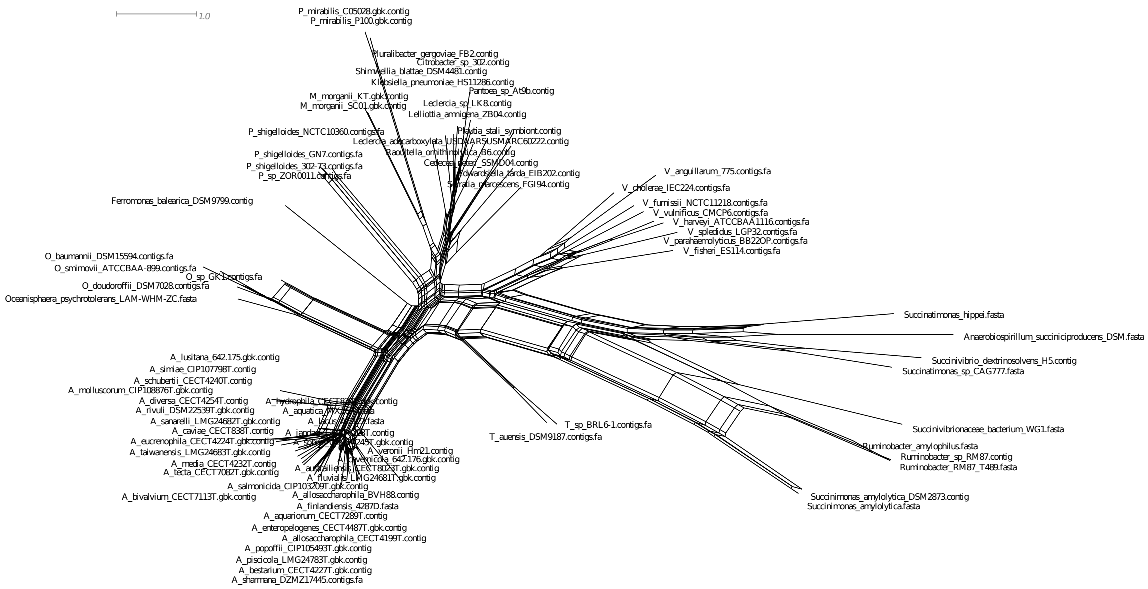


**Fig. S6.** Splits graph diagram for the Aeromonadales dataset built using Splitstree4. The tANI method served provided the distance matrix used to build the graph. The graph further backs the notion that the Aeromonadacea likely need to be revised, as the Succinivibrionaceae place further from the Aeromonas than those of the Enterobacteria.

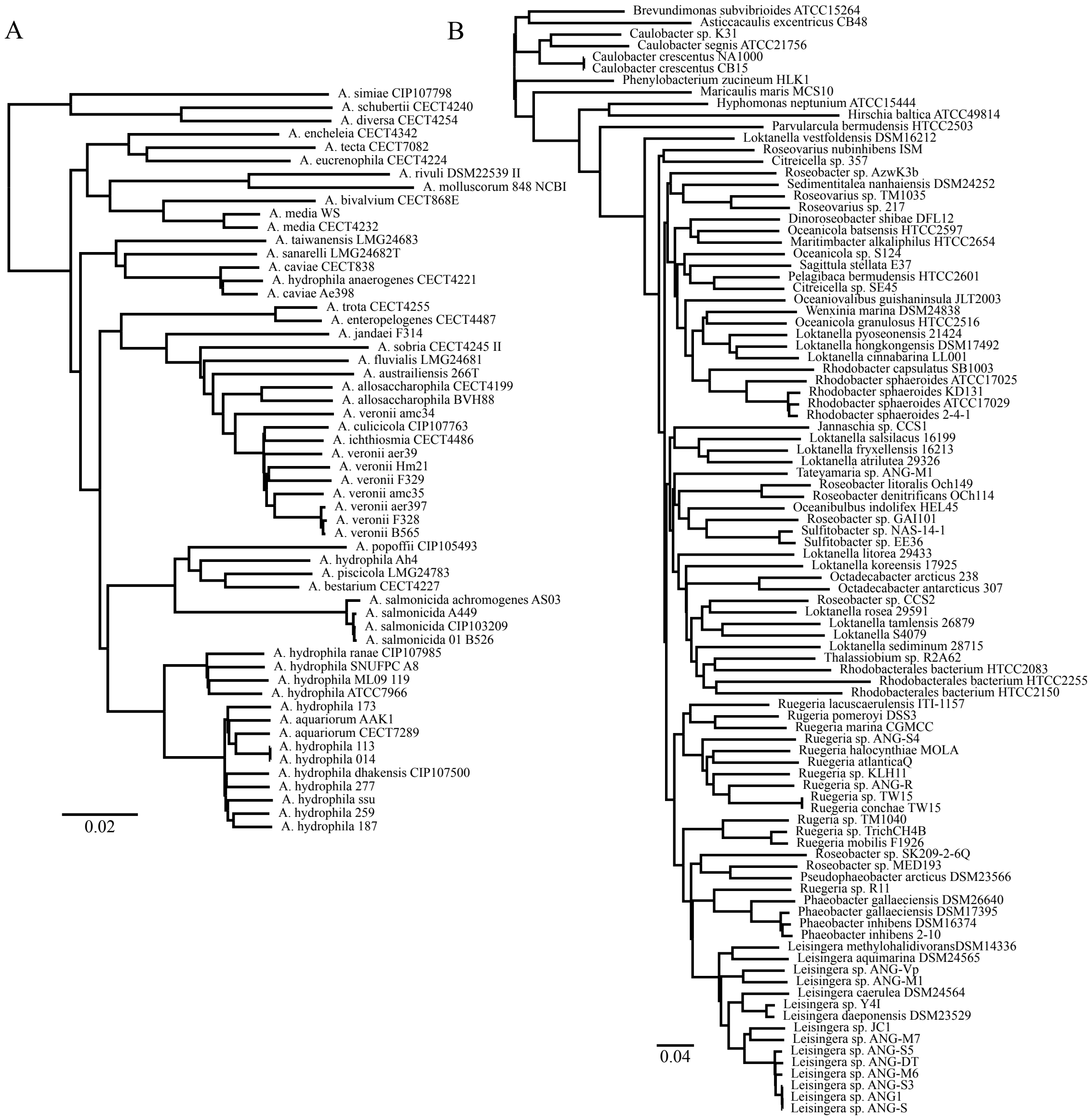


**Fig. S7.** Phylogenies derived from another whole-genome method, MashTree (37). (A) is constructed from the Aeromonas dataset, and (B) is built from the Rhodobacterales dataset.

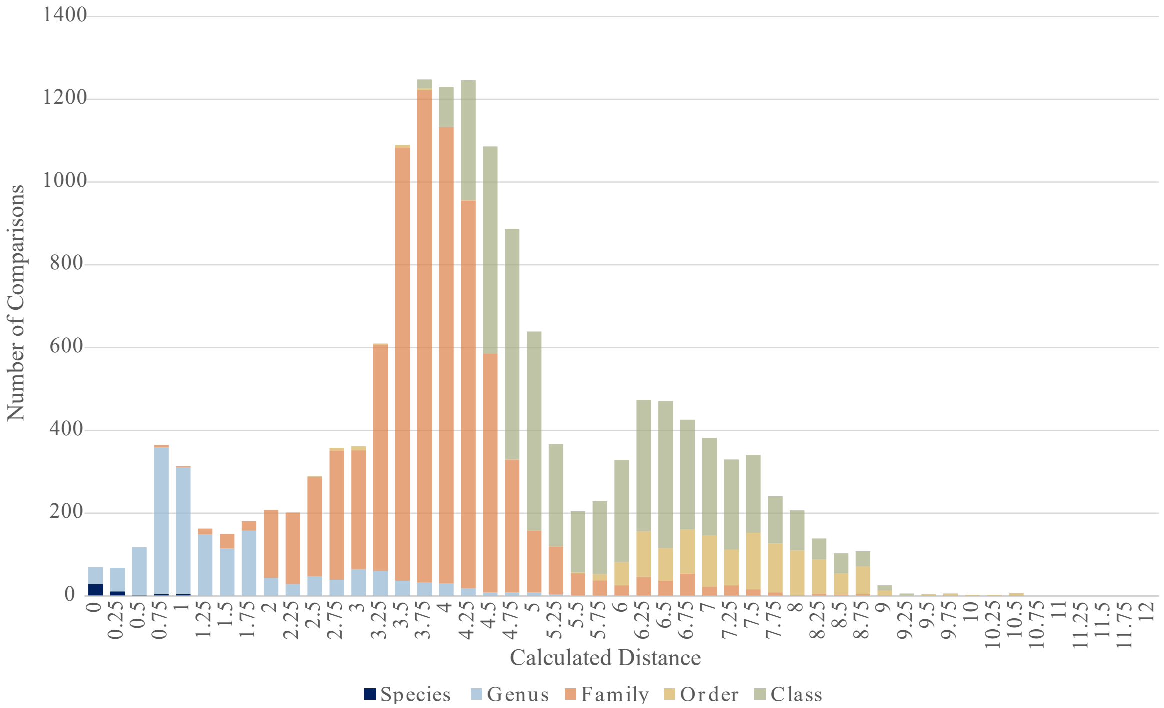


**Fig. S8.** Histograms of tANI values for taxonomic rank comparisons in our datasets using the uncorrected taxonomy as derived from NCBI. Compare Fig. 7 for a similar histogram using the data after correction for misclassified taxa.

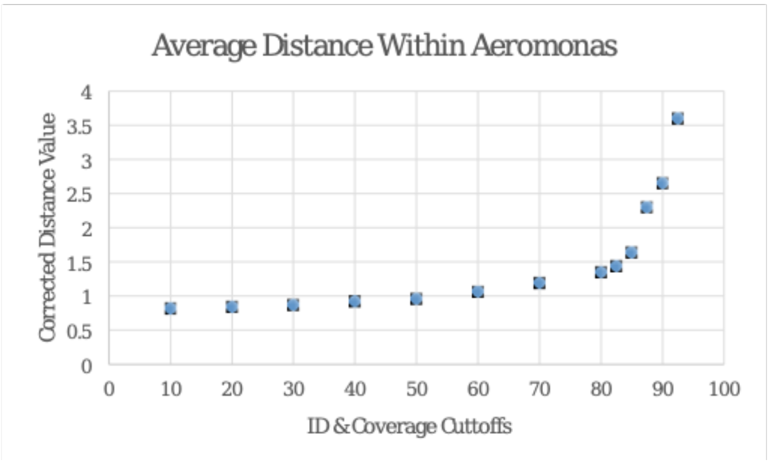


**Fig. S9.** Plot of the average distance of genome-genome tANI calculations within the Aeromonas dataset using varied percent identity and coverage cutoffs. This provided context to which regions of the plot would be best while building phylogenies. Specifically those cutoffs at which the distance was not saturated as a result of including low quality information (10-60), and not inflating as a result of filtering too much information (80+).

**Supplemental Table 1: Full Dataset Description**

| Dataset | Species Name | Strain | Genome Length (Mbp) | No. of Scaffolds | Accession Number |
|---|---|---|---|---|---|
| **Aeromonas** | *A allosaccharophila* | BVH88 | 4.71 | 131 | NZ_CDCB000000000.1 |
| | *A allosaccharophila* | CECT4199 | 4.66 | 120 | NZ_CDBR000000000.1 |
| | *A austrailiensis* | 266T | 4.11 | 113 | NZ_CDDH000000000.1 |
| | *A bestarium* | CECT4227 | 4.69 | 41 | NZ_CDDA000000000.1 |
| | *A bivalvium* | CECT868E | 5.50 | 1112 | NZ_CDBT000000000.1 |
| | *A caviae* | Ae398 | 4.44 | 149 | NZ_CACP000000000.1 |
| | *A caviae* | CECT838 | 4.47 | 111 | NZ_CDBK000000000.1 |
| | *A caviae* | CECT4221 | 4.58 | 332 | NZ_CDBS000000000.1 |
| | *A dhakensis* | AAK1 | 4.76 | 36 | NZ_BAFL000000000.1 |
| | *A dhakensis* | CECT7289 | 4.69 | 78 | NZ_CDBP000000000.1 |
| | *A dhakensis* | 116 | 4.68 | 45 | NZ_ANPN000000000.1 |
| | *A dhakensis* | 187 | 4.78 | 59 | NZ_AOBO000000000.1 |
| | *A dhakensis* | CIP107500 | 4.71 | 73 | NZ_CDBH000000000.1 |
| | *A diversa* | CECT4254 | 4.06 | 37 | NZ_CDCE000000000.1 |
| | *A encheleia* | CECT4342 | 4.47 | 35 | NZ_CDDI000000000.1 |
| | *A enteropelogenes* | CECT4487 | 4.47 | 46 | NZ_CDCG000000000.1 |
| | *A enteropelogenes* | CECT4255 | 4.34 | 27 | NZ_CDDE000000000.1 |
| | *A eucrenophila* | CECT4224 | 4.54 | 22 | NZ_CDDF000000000.1 |
| | *A fluvialis* | LMG24681 | 3.90 | 76 | NZ_CDBO000000000.1 |
| | *A hydrophila* | 14 | 4.67 | 75 | NZ_AOBM000000000.1 |
| | *A hydrophila* | 173 | 4.79 | 74 | NZ_AOBN000000000.1 |
| | *A hydrophila* | 259 | 4.70 | 80 | NZ_AOBP000000000.1 |
| | *A hydrophila* | 277 | 4.79 | 41 | NZ_AOBQ000000000.1 |
| | *A hydrophila* | ATCC7966 | 4.74 | 1 | NC_0085700.1 |
| | *A hydrophila* | ML09 119 | 5.02 | 1 | NC_0212900.1 |
| | *A hydrophila* | SNUFPC-A8 | 4.97 | 41 | NZ_AMQA000000000.1 |
| | *A hydrophila* | ssu | 4.94 | 2 | NZ_AGWR000000000.1 |
| | *A hydrophila subsp. Ranae* | CIP107985 | 4.68 | 107 | NZ_CDDC000000000.1 |
| | *A jandaei* | CECT 4228 | 4.50 | 58 | NZ_CDBV000000000.1 |
| | *A media* | CECT4232 | 4.48 | 233 | NZ_CDBZ000000000.1 |
| | *A media* | WS | 4.32 | 258 | NZ_CP0075670.1 |
| | *A molluscorum* | 848 | 4.24 | 309 | NZ_AQGQ000000000.1 |
| | *A piscicola* | LMG 24783 | 5.18 | 91 | NZ_CDBL000000000.1 |
| | *A popoffii* | CIP105493 | 4.76 | 105 | NZ_CDBI000000000.1 |
| | *A rivuli* | DSM22539 II | 4.53 | 102 | NZ_CDBJ000000000.1 |
| | *A salmonicida* | 01 B526 | 4.93 | 604 | NZ_AGVO000000000.1 |
| | *A salmonicida* | A449 | 5.04 | 6 | NC_0093480.1 |
| | *A salmonicida* | CIP103209 | 4.74 | 128 | NZ_CDDW000000000.1 |
| | *A salmonicida subsp. Achromogenes* | AS03 | 4.45 | 342 | NZ_AMQG000000000.1 |
| | *A sanarelli* | LMG24682T | 4.19 | 98 | NZ_CDBN000000000.1 |
| | *A schubertii* | CECT4240 | 4.13 | 111 | NZ_CDDB000000000.1 |
| | *A simiae* | CIP107798 | 3.99 | 100 | NZ_CDBY000000000.1 |
| | *A sobria* | CECT4245 | 4.68 | 48 | NZ_CDBW000000000.1 |
| | *A sp. nov* | Ah4 | 4.87 | 41 | SAMEA2752429 |
| | *A sp. nov* | amc34 | 4.58 | 1 | NZ_AGWU000000000.1 |
| | *A taiwanensis* | LMG24683 | 5.08 | 987 | NZ_CDDD000000000.1 |
| | *A tecta* | CECT7082 | 4.76 | 51 | NZ_CDCA000000000.1 |
| | *A veronii* | CIP107763 | 4.43 | 64 | NZ_CDDU000000000.1 |
| | *A veronii* | CECT4486 | 4.41 | 66 | NZ_CDBU000000000.1 |
| | *A veronii* | aer39 | 4.42 | 4 | NZ_AGWT000000000.1 |
| | *A veronii* | aer397 | 4.50 | 5 | NZ_AGWV000000000.1 |

| | | | | | |
|---|---|---|---|---|---|
| | *A veronii* | amc35 | 4.57 | 2 | NZ_AGWW000000000.1 |
| | *A veronii* | B565 | 4.55 | 1 | NC_0154240.1 |
| | *A veronii* | F328 | 4.52 | 52 | NZ_CDDK000000000.1 |
| | *A veronii* | Hm21 | 4.68 | 50 | NZ_ATFB000000000.1 |
| ***Aeromonadales*** | *A allosaccharophila* | BVH88 | 4.71 | 131 | NZ_CDCB000000000.1 |
| | *A allosaccharophila* | CECT4199 | 4.66 | 120 | NZ_CDBR000000000.1 |
| | *A aquariorum* | CECT7289T | 4.69 | 78 | NZ_CDBP00000000.1 |
| | *A aquatica* | MX16A | 4.78 | 1 | NZ_CP018201.1 |
| | *A austrailiensis* | 266T | 4.11 | 113 | NZ_CDDH000000000.1 |
| | *A bestarium* | CECT4227 | 4.69 | 41 | NZ_CDDA000000000.1 |
| | *A bivalvium* | CECT868E | 5.50 | 1112 | NZ_CDBT000000000.1 |
| | *A cavernicola* | 642.176 | 3.92 | 341 | NZ_PGGC00000000.1 |
| | *A caviae* | CECT838T | 4.47 | 111 | NZ_CDBK000000000.1 |
| | *A diversa* | CECT4254 | 4.06 | 37 | NZ_CDCE000000000.1 |
| | *A enteropelogenes* | CECT4487 | 4.47 | 46 | NZ_CDCG000000000.1 |
| | *A eucrenophila* | CECT4224 | 4.54 | 22 | NZ_CDDF000000000.1 |
| | *A finlandiensis* | 4287D | 4.72 | 376 | JRGK01000001.1 |
| | *A fluvialis* | LMG24681 | 3.90 | 76 | NZ_CDBO000000000.1 |
| | *A hydrophila* | CECT839T | 4.74 | 1 | NC_0085700.1 |
| | *A jandaei* | CECT 4228 | 4.50 | 58 | NZ_CDBV000000000.1 |
| | *A lacus* | AE122 | 4.39 | 196 | JRGM01000001.1 |
| | *A lusitana* | 642.175 | 4.55 | 67 | PGCP00000000.1 |
| | *A media* | CECT4232 | 4.48 | 233 | NZ_CDBZ000000000.1 |
| | *A molluscorum* | CIP108876T | 4.24 | 309 | NZ_AQGQ000000000.1 |
| | *A piscicola* | LMG 24783 | 5.18 | 91 | NZ_CDBL000000000.1 |
| | *A popoffii* | CIP105493 | 4.76 | 105 | NZ_CDBI000000000.1 |
| | *A rivuli* | DSM22539 II | 4.53 | 102 | NZ_CDBJ000000000.1 |
| | *A salmonicida* | CIP103209 | 4.74 | 128 | NZ_CDDW000000000.1 |
| | *A sanarelli* | LMG24682T | 4.19 | 98 | NZ_CDBN000000000.1 |
| | *A schubertii* | CECT4240 | 4.13 | 111 | NZ_CDDB000000000.1 |
| | *A simiae* | CIP107798 | 3.99 | 100 | NZ_CDBY000000000.1 |
| | *A sobria* | CECT4245 | 4.68 | 48 | NZ_CDBW000000000.1 |
| | *A taiwanensis* | LMG24683 | 5.08 | 987 | NZ_CDDD000000000.1 |
| | *A tecta* | CECT7082 | 4.76 | 51 | NZ_CDCA000000000.1 |
| | *A veronii* | Hm21 | 4.68 | 50 | NZ_ATFB000000000.1 |
| | *Anaerobiospirillum succiniciproducens* | DSM | 3.80 | 155 | AXWV01000001.1 |
| | *Cedecea neteri* | SSMD04 | 4.88 | 1 | NZ_CP009451.1 |
| | *Citrobacter* | sp 302 | 5.02 | 9 | NZ_KI391984.1 |
| | *Edwardsiella tarda* | EIB202 | 3.80 | 2 | NC_013508.1 |
| | *Ferromonas balearica* | DSM9799 | 4.28 | 1 | NC_014541.1 |
| | *Klebsiella pneumoniae* | HS11286 | 5.68 | 7 | NC_016845.1 |
| | *Leclercia adecarboxylata* | USDAARSUSMARC60222 | 4.80 | 1 | NZ_CP013990.1 |
| | *Leclercia* | sp LK8 | 5.21 | 75 | NZ_LDUO01000001.1 |
| | *Lelliottia amnigena* | ZB04 | 4.62 | 1 | NZ_CP015774.1 |
| | *Morganella morganii* | KT | 3.83 | 58 | NC_020418.1 |
| | *Morganella morganii* | SC01 | 4.15 | 63 | NZ_AMWL00000000.2 |
| | *Oceanimonas baumannii* | DSM15594s | 3.75 | 31 | 2593339295* |
| | *Oceanimonas doudoroffii* | DSM7028s | 3.83 | 194 | 2506520053* |
| | *Oceanimonas smirnovii* | ATCCBAA-899s | 3.28 | 28 | NZ_ARMW00000000.1 |
| | *Oceanimonas* | sp GK1s | 3.51 | 1 | NC_016745.1 |
| | *Oceanisphaera psychrotolerans* | LAM-WHM-ZC | 3.82 | 70 | MDKE01000001.1 |
| | *Plesiomonas shigelloides* | 302-73s | 3.91 | 389 | NZ_AQQO00000000.1 |
| | *Plesiomonas shigelloides* | GN7s | 3.92 | 83 | NZ_JWHQ00000000.1 |
| | *Plesiomonas shigelloides* | NCTC10360s | 2.46 | 1 | NZ_LT575468.1 |
| | *Plesiomonas* | sp ZOR0011s | 3.84 | 152 | NZ_JRKB00000000.1 |
| | *Proteus mirabilis* | C05028 | 3.79 | 85 | NZ_ANBT00000000.1 |

| | | | | | |
|---|---|---|---|---|---|
| | *Proteus mirabilis* | P100 | 4.15 | 126 | 2740892266* |
| | *Pantoea* | sp At9b | 6.31 | 6 | NC_014837.1 |
| | *Plautia* | stali symbiont | 4.09 | 3 | NC_022546.1 |
| | *Pluralibacter gergoviae* | FB2 | 5.49 | 1 | NZ_CP009450.1 |
| | *Raoultella ornithinolytica* | B6 | 5.40 | 1 | NC_021066.1 |
| | *Ruminobacter amylophilus* | DSM 1361 | 2.82 | 117 | FOXF01000116.1 |
| | *Ruminobacter* | RM87 T489 | 2.86 | 123 | JNKD01000001.1 |
| | *Ruminobacter* | sp RM87 | 2.86 | 123 | NZ_JNKD01000001.1 |
| | *Serratia marcescens* | FGI94 | 4.86 | 1 | NC_020064.1 |
| | *Shimwellia blattae* | DSM4481 | 4.16 | 1 | NC_017910.1 |
| | *Succinimonas amylolytica* | DSM 2873 | 3.96 | 23 | KB899636.1 |
| | *Succinatimonas hippei* | YIT 12066 | 2.31 | 141 | GL830939.1 |
| | *Succinatimonas* | sp CAG777 | 2.25 | 71 | HF987897.1 |
| | *Succinivibrionaceae bacterium* | WG1 | 2.95 | 43 | GL995195.1 |
| | *Succinimonas amylolytica* | DSM2873 | 3.96 | 23 | NZ_KB899636.1 |
| | *Succinivibrio dextrinosolvens* | H5 | 2.68 | 106 | NZ_KL370853.1 |
| | *Tolumonas auensis* | DSM9187s | 3.47 | 1 | NC_012691.1 |
| | *Tolumonas* | sp BRL6-1s | 3.63 | 9 | NZ_AZUK00000000.1 |
| | *Vibrio anguillarum* | 775s | 4.05 | 2 | NC_015633.1 |
| | *Vibrio cholerae* | IEC224s | 4.08 | 2 | NC_016944.1 |
| | *Vibrio fisheri* | ES114s | 4.27 | 3 | NC_006840.2 |
| | *Vibrio furnissii* | NCTC11218s | 4.92 | 2 | NC_016602.1 |
| | *Vibrio harveyi* | ATCCBAA1116s | 6.06 | 3 | NC_009777.1 |
| | *Vibrio parahaemolyticus* | BB22OPs | 5.10 | 2 | NC_019955.1 |
| | *Vibrio spledidus* | LGP32s | 4.97 | 2 | NC_011744.2 |
| | *Vibrio vulnificus* | CMCP6s | 5.13 | 2 | NC_004459.3 |
| | *Asticcacaulis excentricus* | CB48 | 4.31 | 4 | NC_014817.1 |
| | *Brevundimonas subvibrioides* | ATCC15264 | 3.45 | 1 | NC_014375.1 |
| | *Caulobacter* | K31 | 5.89 | 3 | NC_002696.2 |
| | *Caulobacter crescentus* | CB15 | 4.02 | 1 | NC_011916.1 |
| | *Caulobacter crescentus* | NA1000 | 4.04 | 1 | NC_014100.1 |
| | *Caulobacter segnis* | ATCC21756 | 4.66 | 1 | NC_010333.1 |
| | *Citreicella* | 357 | 4.60 | 180 | NZ_AJKJ01000164.1 |
| | *Citreicella* | SE45 | 5.52 | 9 | NZ_GG704601.1 |
| | *Dinoroseobacter shibae* | DFL12 | 4.42 | 6 | NC_009959.1 |
| | *Hirschia baltica* | ATCC49814 | 3.54 | 2 | NC_012983.1 |
| | *Hyphomonas neptunium* | ATCC15444 | 3.71 | 1 | NC_008358.1 |
| | *Jannaschia* | CCS1 | 4.40 | 2 | NC_007802.1 |
| | *Leisingera* | ANG-DT | 4.60 | 91 | NZ_JWLE00000000.1 |
| | *Leisingera* | ANG-M1 | 5.38 | 92 | NZ_JWLC00000000.1 |
| | *Leisingera* | ANG-M6 | 4.54 | 54 | NZ_JWLG00000000.1 |
| | *Leisingera* | ANG-M7 | 4.58 | 58 | NC_023146.1 |
| | *Leisingera* | ANG-S | 4.57 | 68 | NZ_JWLM00000000.1 |
| | *Leisingera* | ANG-S3 | 4.60 | 70 | NZ_JWLF00000000.1 |
| | *Leisingera* | ANG-S5 | 4.66 | 43 | NZ_JWLH00000000.1 |
| | *Leisingera* | ANG-Vp | 5.15 | 143 | NZ_JWLD00000000.1 |
| | *Leisingera* | ANG1 | 4.60 | 26 | NZ_AFCF00000000.2 |
| | *Leisingera* | Y4I | 4.34 | 5 | NZ_DS995283.1 |
| | *Leisingera* | JC1 | 5.19 | 168 | NZ_LYUZ00000000.1 |
| | *Leisingera aquimarina* | DSM24565 | 5.34 | 15 | NC_023146.1 |
| | *Leisingera caerulea* | DSM24564 | 5.34 | 21 | NZ_AXBI00000000.1 |
| | *Leisingera daeponensis* | DSM23529 | 4.64 | 12 | NZ_AXBD00000000.1 |
| | *Leisingera methylohalidivorans* | DSM14336 | 4.65 | 3 | NZ_DS995283.1 |
| | *Loktanella* | S4079 | 3.56 | 43 | NZ_JXYE00000000.1 |
| | *Loktanella atrilutea* | 29326 | 4.21 | 46 | NZ_FQUE00000000.1 |
| | *Loktanella cinnabarina* | LL001 | 3.90 | 192 | NZ_BATB00000000.1 |

| | | | | | |
|---|---|---|---|---|---|
| Rhodobacterales | *Loktanella fryxellensis* | 16213 | 3.55 | 75 | FOCI00000000.1 |
| | *Loktanella hongkongensis* | DSM17492 | 3.19 | 16 | NZ_KB823002.1 |
| | *Loktanella koreensis* | 17925 | 3.65 | 4 | FOIZ00000000.1 |
| | *Loktanella litorea* | 29433 | 3.32 | 8 | FOZM00000000.1 |
| | *Loktanella pyoseonensis* | 21424 | 3.91 | 22 | NZ_FTPR00000000.1 |
| | *Loktanella rosea* | 29591 | 3.51 | 5 | FNAT00000000.1 |
| | *Loktanella salsilacus* | 16199 | 4.13 | 77 | FOTF00000000.1 |
| | *Loktanella sediminum* | 28715 | 3.26 | 16 | NZ_FQXB00000000.1 |
| | *Loktanella tamlensis* | 26879 | 3.19 | 9 | FOYP00000000.1 |
| | *Loktanella vestfoldensis* | DSM16212 | 3.72 | 49 | NZ_ARNL00000000.1 |
| | *Maricaulis maris* | MCS10 | 3.37 | 1 | NC_008347.1 |
| | *Maritimbacter alkaliphilus* | HTCC2654 | 4.54 | 7 | NZ_AAMT01000046.1 |
| | *Oceanibulbus indolifex* | HEL45 | 4.11 | 105 | NZ_ABID01000017.1 |
| | *Oceanicola* | S124 | 4.65 | 339 | NZ_AAMO01000007.1 |
| | *Oceanicola batsensis* | HTCC2597 | 4.44 | 7 | NZ_CH724110.1 |
| | *Oceanicola granulosus* | HTCC2516 | 4.05 | 9 | NZ_AFPM01000263.1 |
| | *Oceaniovalibus guishaninsula* | JLT2003 | 2.90 | 68 | NZ_AMGO01000046.1 |
| | *Octadecabacter antarcticus* | 307 | 4.88 | 2 | NC_020911.1 |
| | *Octadecabacter arcticus* | 238 | 5.48 | 3 | NC_020909.1 |
| | *Parvularcula bermudensis* | HTCC2503 | 2.90 | 1 | NC_014414.1 |
| | *Pelagibaca bermudensis* | HTCC2601 | 5.48 | 6 | NZ_DS022279.1 |
| | *Phaeobacter gallaeciensis* | DSM17395 | 4.23 | 4 | NC_018290.1 |
| | *Phaeobacter gallaeciensis* | DSM26640 | 4.54 | 8 | NC_023143.1 |
| | *Phaeobacter inhibens* | 2-10 | 4.16 | 4 | NC_018423.1 |
| | *Phaeobacter inhibens* | DSM16374 | 4.13 | 8 | NZ_AXBB00000000.1 |
| | *Phenylobacterium zucineum* | HLK1 | 4.38 | 2 | NC_011143.1 |
| | *Pseudophaeobacter arcticus* | DSM23566 | 5.05 | 8 | NZ_AXBF00000000.1 |
| | *Rhodobacter capsulatus* | SB1003 | 3.87 | 2 | NC_014035.1 |
| | *Rhodobacter haeroides* | 2-4-1 | 4.60 | 7 | NC_009007.1 |
| | *Rhodobacter haeroides* | ATCC17025 | 4.56 | 6 | NC_009430.1 |
| | *Rhodobacter haeroides* | ATCC17029 | 4.49 | 3 | NC_009049.1 |
| | *Rhodobacter haeroides* | KD131 | 4.71 | 4 | NC_011960.1 |
| | *Rhodobacterales bacterium* | HTCC2083 | 4.02 | 5 | NZ_DS995280.1 |
| | *Rhodobacterales bacterium* | HTCC2150 | 3.58 | 25 | NZ_AAXZ01000012.1 |
| | *Rhodobacterales bacterium* | HTCC2255 | 2.30 | 2 | NZ_DS022282.1 |
| | *Roseobacter* | AzwK3b | 4.18 | 31 | NC_008388.1 |
| | *Roseobacter* | CCS2 | 3.50 | 11 | NC_015730.1 |
| | *Roseobacter* | GAI101 | 4.53 | 9 | NZ_ABCR01000029.1 |
| | *Roseobacter* | MED193 | 4.67 | 4 | NZ_AAYB01000010.1 |
| | *Roseobacter* | SK209-2-6 | 4.56 | 29 | NZ_DS999219.1 |
| | *Roseobacter denitrificans* | OCh114 | 4.33 | 5 | NZ_AANB01000005.1 |
| | *Roseobacter litoralis* | Och149 | 4.75 | 4 | NZ_AAYC01000028.1 |
| | *Roseovarius* | 217 | 4.77 | 6 | NZ_CH724156.1 |
| | *Roseovarius* | TM1035 | 4.21 | 15 | NZ_CH902585.1 |
| | *Roseovarius nubinhibens* | ISM | 3.68 | 4 | NZ_ABCL01000004.1 |
| | *Ruegeria* | ANG-R | 4.68 | 41 | NZ_JWLJ00000000.1 |
| | *Ruegeria* | ANG-S4 | 4.54 | 20 | NZ_JWLK00000000.1 |
| | *Ruegeria* | KLH11 | 4.49 | 6 | NZ_DS999534.1 |
| | *Ruegeria* | R11 | 3.82 | 2 | NZ_DS999534.1 |
| | *Ruegeria* | TrichCH4B | 4.67 | 129 | NZ_DS999055.1 |
| | *Ruegeria* | TW15 | 4.49 | 28 | NZ_AEYW01000007.1 |
| | *Ruegeria* | TM1040 | 4.15 | 3 | NC_006569.1 |
| | *Ruegeria atlantica* | CECT4293 | 4.82 | 67 | NZ_CYPS00000000.1 |
| | *Ruegeria conchae* | TW15 | 4.49 | 28 | NZ_AEYW00000000.1 |
| | *Ruegeria halocynthiae* | MOLA | 4.31 | 19 | NZ_JQEZ00000000.1 |
| | *Ruegeria lacuscaerulensis* | ITI-1157 | 3.52 | 47 | NZ_FQYJ00000000.1 |

| | | | | | |
|---|---|---|---|---|---|
| | *Ruegeria marina* | CGMCC | 5.00 | 53 | FMZV00000000.1 |
| | *Ruegeria mobilis* | F1926 | 4.83 | 5 | NZ_CP015230.1 |
| | *Rugeria pomeroyi* | DSS3 | 4.60 | 2 | NC_008043.1 |
| | *Sagittula stellata* | E37 | 5.26 | 39 | NZ_AAYA01000035.1 |
| | *Sedimentitalea nanhaiensis* | DSM24252 | 4.95 | 30 | NZ_AXBG00000000 |
| | *Sulfitobacter* | EE36 | 3.37 | 4 | NZ_CH959310.1 |
| | *Sulfitobacter* | NAS-14-1 | 4.01 | 11 | NZ_CH959313.1 |
| | *Tateyamaria* | ANG-M1 | 4.43 | 32 | NZ_JWLL00000000 |
| | *Thalassiobium* | R2A62 | 3.49 | 1 | NZ_GG697169.2 |
| | *Wenxinia marina* | DSM24838 | 4.18 | 41 | NZ_KB902299.1 |
| ***Frankiales*** | *Cryptosporangium arvum* | 44712 | 9.20 | 1 | NZ_JFBT00000000.1 |
| | *Cryptosporangium aurantiacum* | DSM46144 | 9.58 | 44 | NZ_FRCS00000000.1 |
| | *Frankia* | CcI6 | 5.58 | 136 | GCA_000503735.2 |
| | *Frankia* | 45899 | 9.54 | 83 | GCA_001536285.1 |
| | *Frankia* | ACN1ag | 7.52 | 90 | GCA_001414035.1 |
| | *Frankia* | Allo2 | 5.35 | 110 | GCA_000733325.1 |
| | *Frankia* | AvcI1 | 7.74 | 77 | GCA_001420875.1 |
| | *Frankia* | BMG523 | 5.27 | 166 | GCA_000685765.2 |
| | *Frankia* | BMG530 | 5.82 | 95 | GCA_001983005.1 |
| | *Frankia* | BMG536 | 11.20 | 280 | GCA_001854805.1 |
| | *Frankia* | BRAAY231001 | 5.23 | 180 | GCA_001636575.1 |
| | *Frankia* | Cc117 | 8.36 | 195 | GCA_001854655.1 |
| | *Frankia* | CcI156 | 5.33 | 145 | GCA_001983015.1 |
| | *Frankia* | CcI49 | 9.76 | 78 | GCA_001983215.1 |
| | *Frankia* | CED | 5.00 | 120 | GCA_000732115.1 |
| | *Frankia* | CgIM4 | 5.20 | 135 | GCA_001756285.1 |
| | *Frankia* | CgIS1 | 8.03 | 289 | GCA_001854725.1 |
| | *Frankia* | CN3FCB1 | 9.98 | 2 | GCA_000235425.3 |
| | *Frankia coriariae* | BMG51 | 5.80 | 116 | NZ_JWIO00000000.1 |
| | *Frankia* | CpI1PFF861001 | 7.62 | 143 | GCA_001421075.1 |
| | *Frankia* | CpI1SFF36 | 7.62 | 153 | GCA_000948395.1 |
| | *Frankia* | DC12 | 6.88 | 1 | GCA_000966285.1 |
| | *Frankia* | Dg2 | 5.90 | 2738 | GCA_900067225.1 |
| | *Frankia discariae* | BCU110501 | 7.89 | 194 | NZ_ARDT00000000.1 |
| | *Frankia* | EAN1pec | 8.98 | 1 | NC_009921.1 |
| | *Frankia* | EI5cUG551001 | 6.62 | 159 | GCA_001636565.1 |
| | *Frankia elaeagni* | BMG512 | 7.59 | 135 | NZ_ARFH00000000.1 |
| | *Frankia inefficax* | EuI1c | 8.82 | 1 | NC_014666.1 |
| | *Frankia* | EUN1fctg00163 | 9.35 | 396 | GCA_000177675.1 |
| | *Frankia* | EUN1h | 9.91 | 129 | GCA_001854645.1 |
| | *Frankia* | Iso899 | 5.10 | 67 | GCA_000421445.1 |
| | *Frankia* | NRRLB16219 | 5.26 | 135 | GCA_001854695.1 |
| | *Frankia asymbiotica* | NRRLB16386 | 9.44 | 174 | NZ_MOMC00000000.1 |
| | *Frankia* | QA3 | 7.59 | 1 | NZ_CM001489.1 |
| | *Frankia* | R43 | 10.45 | 46 | GCA_001306465.1 |
| | *Frankia* | Symbiont of Datisca glomerata | 5.34 | 3 | NC_015656.1 |
| | *Frankia* | Thr | 5.31 | 169 | GCA_000611815.2 |
| | *Jatrophihabitans endophyticus* | 45627 | 4.48 | 10 | NZ_FQVU00000000.1 |
| | *Sporichthya polymorpha* | 43042 | 5.50 | 1 | NZ_AQZX00000000.1 |
| | *Frankia alni* | ACN14A | 7.50 | 1 | NC_008278.1 |
| | *Frankia casuarinae* | CcI3 | 5.43 | 1 | NC_007777.1 |
| | *Frankineae bacterium* | MT45 | 4.23 | 1 | GCA_900100325.1 |

**Supplemental Table 2.**

Tree Certaintly Averages comparing  support sets to the best trees using different approaches.

| Dataset | Tree | Bootstrap Set | Avg. IC |
|---|---|---|---|
| Aeromonas | tANI | tANI | 0.86 |
| | tANI | MLSA | 0.33 |
| | tANI | Core | 0.61 |
| | MLSA | tANI | 0.27 |
| | MLSA | MLSA | 0.65 |
| | MLSA | Core | 0.24 |
| | Core | tANI | 0.61 |
| | Core | MLSA | 0.37 |
| | Core | Core | 0.87 |
| | Mashtree | tANI | 0.47 |
| | Mashtree | MLSA | 0.28 |
| | Mashtree | Core | 0.47 |
| Rhodobacterales | tANI | tANI | 0.80 |
| | tANI | MLSA | 0.39 |
| | MLSA | MLSA | 0.80 |
| | MLSA | tANI | 0.37 |
| | Mashtree | MLSA | 0.25 |
| | Mashtree | tANI | 0.23 |

**Supplemental Table 3.** Changes for Species Level Cutoff Comparison

| **Current ID** | **ID Used in Cutoff Comparison*** |
|---|---|
| *Leisingera* daeponensis DSM23529 | Leisingera_2 |
| *Leisingera* sp ANG1 | Leisingera_1 |
| *Leisingera* sp ANG-M6 | Leisingera_1 |
| *Leisingera* sp ANG-S | Leisingera_1 |
| *Leisingera* sp ANG-S3 | Leisingera_1 |
| *Leisingera* sp ANG-S5 | Leisingera_1 |
| *Leisingera* sp Y4I | Leisingera_2 |
| *Leisingera* sp ANG-DT | Leisingera_1 |
| *Sulfitobacter* sp EE36 | Sulfitobacter_1 |
| *Sulfitobacter* sp NAS-14-1 | Sulfitobacter_1 |
| *Ruegeria* conchae TW15 | Ruegeria_conchae_TW15 |
| *Ruegeria* lacuscaerulensis ITI-1157 | Ruegeria_lacusaerulensis |
| *Ruegeria* mobilis F1926 | Ruegeria_mobilis |
| *Ruegeria* sp TrichCH4B | Ruegeria_mobilis |
| *Ruegeria* sp TW15 | Ruegeria_conchae_TW15 |
| *Silicibacter* lacuscaerulensis ITI1157 | Ruegeria_lacusaerulensis |

*This is based off of the analysis of tANI and MLSA phylogenies.

**Supplemental Table 4.** Suggested Reclassifications

| **Current Classification** | **Evidence** | **Recommendation*** |
|---|---|---|
| *Rugeria* sp. TM1040 | Recent studies on similar datasets have revealed findings which closely resemble our own; that being that the group is polyphyletic in nature. These three strains in particular classify outside the main genus in our analyses. | Further investigate the relationship of these three strains to the rest of *Ruegeria*. They may need to be classified as a separate genus. Additionally, reclassify TrichCH4B as a mobilis strain. |
| *Ruegeria* mobilis F1926 | | |
| *Ruegeria* sp. TrichCH4B | | |
| *Loktanella* cinnabarina LL001 | Identical reasons to those stated for the *Ruegeria* group. | Investigate the relationship of these species to the rest of the genus. They likely need to be classified as a separate genus |
| *Loktanella* hongkongensis DSM17492 | | |
| *Loktanella* pyoseonensis 21424 | | |
| *Ruegeria* sp. R11 | Consistently groups as an out-group to Phaeobacter no matter the methodology | Investigate relationship to Phaeobacter, and closely related taxa. Likely not a member of *Ruegeria* |
| Anaerobiospirillum succiniciproducens DSM | The classification of this grouping as part of Aeromonadales is based off a 1999 study by Hippe et al., which appears to be the primary source of phylogenetic analysis for the classification. However, this study had did not have the sampling now available, and there were no further phylogenies to support the claim. Current sampling and phylogenies indicate that this group is further away from the rest of the Aeromonadales than Enterobacteraceae and members of Oceanismonas. | The family Succinivibrionaceae likely does not belong within Aeromonadales. However, due to this group being plagued by long branches, more sampling will be needed to understand the true nature of this group, and how the members within are related to one another. |
| Oceanisphaera psychrotolerans LAM-WHM-ZC | | |
| Ruminobacter amylophilus DSM 1361 | | |
| Ruminobacter RM87 T489 | | |
| Succinatimonas hippei YIT 12066 | | |
| Succinatimonas sp CAG777 | | |
| Succinimonas amylolytica DSM2873 | | |
| Succinivibrio dextrinosolvens H5 | | |
| Succinivibrio Phil9 | | |
| Succinivibrionaceae bacterium WG1 | | |

***These suggested changes were applied to classifications in the higher order ROC comparisons**

**Supplemental Table 5.**

Frankiales MLSA Genes

| Gene | Location on NC_007777.1 |
|---|---|
| glutamate synthase beta subunit | (3573463-3574905) |
| glutamate synthase alpha subunit | (3574965-3579521) |
| ribosomal protein large subunit 1 | (659953-660669) |
| ribosomal protein large subunit 2 | (679418-680251) |
| ribosomal protein large subunit 3 | (677637-678344) |
| ribosomal protein large subunit 4 | (678341-679078) |
| ribosomal protein small subunit 1 | (1259062-1260540) |
| ribosomal protein small subunit 2 | (4280241-4281110) |
| ribosomal protein small subunit 3 | (681044-681991) |
| ribosomal protein small subunit 4 | (692551-693180) |
| elongation factor Tu | (675819-677012) |
| bipA | (4627929-4629767) |
| ATP synthase subunit alpha | (4442416-4444074) |
| ATP synthase subunit beta | (4439882-4441321) |
| dnaA | (35-1723) |
| dnaK | (5197168-5199018) |
| dnaX | (309543-311981) |
| GAPDH | (1969408-1970415) |
| groEL | (715686-717326) |
| gyrA | (8155-10659) |
| gyrB | (6021-7955) |
| recA | (4210237-4211274) |
| rpoB | (662999-666424) |
| rpoD | (1534111-1535289) |
